# A cellular anatomy of the normal adult human prostate and prostatic urethra

**DOI:** 10.1101/439935

**Authors:** Gervaise H. Henry, Alicia Malewska, Diya B. Joseph, Venkat S. Malladi, Jeon Lee, Jose Torrealba, Ryan J. Mauck, Jeffrey C. Gahan, Ganesh V. Raj, Claus G. Roehrborn, Gary C. Hon, Malcolm P. MacConmara, Jeffrey C. Reese, Ryan C. Hutchinson, Chad M. Vezina, Douglas W. Strand

## Abstract

A cellular anatomy of normal human organs is essential for solving the cellular origins of disease. We report the first comprehensive cellular atlas of the young adult human prostate and prostatic urethra using an iterative process of single cell RNA sequencing and flow cytometry on ~98,000 cells taken from different anatomical regions. Two previously unrecognized epithelial cell types were identified by KRT13 and SCGB1A1 expression and found to be highly similar to hillock and club cells of the proximal lung. It was demonstrated by immunohistochemistry that prostate club and hillock cells are similarly concentrated in the proximal prostate. We also optimized a new flow cytometry antibody panel to improve cell type-specific purification based on newly established cellular markers. The molecular classification, anatomical distribution, and purification methods for each cell type in the human prostate create a powerful new resource for experimental design in human prostate disease.

## Introduction

The design of novel therapies against disease relies on a deep understanding of the identity and function of each cell type within an organ. A three-dimensional cellular anatomy of normal organs is necessary to better understand the processes of age-related repair and disease. These efforts have been largely driven by recent advances in single cell sequencing (to identify cell type) and imaging technologies (to identify cell location). Because of the challenges with procurement of fresh normal human organs and the pronounced anatomical differences between the mouse and human prostate, there remain considerable gaps in our understanding of the functions of specific cell types in prostate disease.

The current zonal anatomy of the human prostate was established by John McNeal using hundreds of cadaver specimens^1^. McNeal’s scheme divides the adult human prostate into an anterior fibromuscular zone and three glandular zones (the central zone surrounds the ejaculatory ducts, the transition zone surrounds the urethra, and the peripheral zone surrounds both). McNeal observed that benign prostatic hyperplasia (BPH) occurs mostly in the transition and central zones, while most prostate cancers are found in the peripheral zone. No study has examined how prostate cell types are distributed across each of McNeal’s zones, a critical step towards identifying the cellular origins of prostate cancer and BPH.

Prostate cell types have been subjectively defined by their shape, gene expression, surface antigens, and relative position in glandular acini^2,3^. These criteria have led to the notion that prostate glands contain three unique epithelial cell types: basal, luminal, and neuroendocrine. Basal epithelia express cytokeratin 5 and the transcription factor p63. Luminal epithelia express cytokeratin 8 and androgen-regulated secretory proteins such as KLK3. A putative intermediate cell ‘state’ between basal and luminal lineages has been defined on the basis of shared expression of basal and luminal cytokeratins^4,5^. Neuroendocrine epithelia express markers such as chromogranin A^6^. Various cell surface antibodies and promoters driving fluorophores in transgenic mice are used to label and isolate basal and luminal epithelia by flow cytometry, but the purity of these putative epithelial cell types has never been evaluated. A lack of established stromal cell type surface markers has completely prevented their identification and isolation.

To properly define human prostate cellular anatomy and create a baseline for understanding the cellular origins of disease, we performed single cell RNA sequencing (scRNA-seq) on ~98,000 cells from five young adult human prostates. This is the first widespread application of scRNA-seq to the non-diseased human prostate. Two new epithelial cell types were identified and new, objective markers were derived for known cell types.

scRNA-seq also revealed flaws in the traditional fluorescence activated cell sorting (FACS) gating strategy for human prostate cell types resulting in contaminated bulk RNA sequencing. Accordingly, we describe an improved purification scheme that includes the ability to purify stromal cell types, which had not been possible. We also used scRNA-seq to identify selective cell markers and performed immunostaining on whole transverse prostate sections to demonstrate regional enrichment of cell types as a means to objectively define prostate zonal anatomy in non-diseased specimens. Given the difficulty of routinely procuring young human prostate specimens, these data provide a valuable resource for establishing a molecular and cellular baseline for understanding changes in human prostate disease.

## Results

### Bulk sequencing of the human prostate cells sorted by FACS suggests impurity

Isolating pure cell populations is critical for functional analysis, yet current prostate cell purification protocols fail to achieve purity because they rely on non-specific definitions of cell identity. For example, most studies use fewer than three cell surface markers to define prostate basal and luminal cells, which forces broad assumptions of cell identity. Some groups employ a pan-epithelial marker (CD326, CD324, or TROP2) with a positive basal marker like CD49f assuming that all CD49f^LO^ epithelia are luminal^7^. Other groups use a combination of positive basal (CD49f) and positive luminal (CD26 or CD38) markers^8,9^. Several other options exist for identifying basal epithelia including Podoplanin, CD104, and CD271^10^. While most labs have historically defined basal epithelia as CD49^fHI^, its tri-modal spectrum of expression (high in basal epithelia and 50% of stroma, low in luminal, and negative in the other 50% of stroma) makes it difficult to objectively establish a negative gate for basal epithelial purity. We previously showed that FACS gating for CD271, CD104, and Podoplanin was superior to CD49f because it establishes a definitive boundary between marker-positive and marker-negative basal epithelia^11^. Regardless of which positive marker is used to identify basal epithelia, a double-negative epithelial gate consistently emerges and has never been characterized. We set out to define this additional epithelial cell gate by comparing its transcriptome to that of basal (CD271^+^) and luminal (CD26^+^) gates.

Benign and malignant prostate diseases are widespread in aging men resulting in a perturbation of cellular transcriptomes. To establish a baseline transcriptome for each cell type, we created a fresh tissue biorepository of prostates from young organ donors aged 18-29. Prostates were dissected and enzymatically dissociated into single cell suspensions (Supplementary Figure 1). Viable cells from each gated epithelial population (basal, luminal, other) were collected via FACS according to our previously published protocol^11^. To determine transcriptomic differences between luminal, basal, and other epithelia gates, cDNA libraries from a bulk population of cells from each FACS gate were prepared for sequencing (Figure 1a). Principal component analysis demonstrates concordance of gated epithelial cells across four normal specimens, a testament to the consistency of our approach (Supplementary Figure 2a).

**Figure 1.**
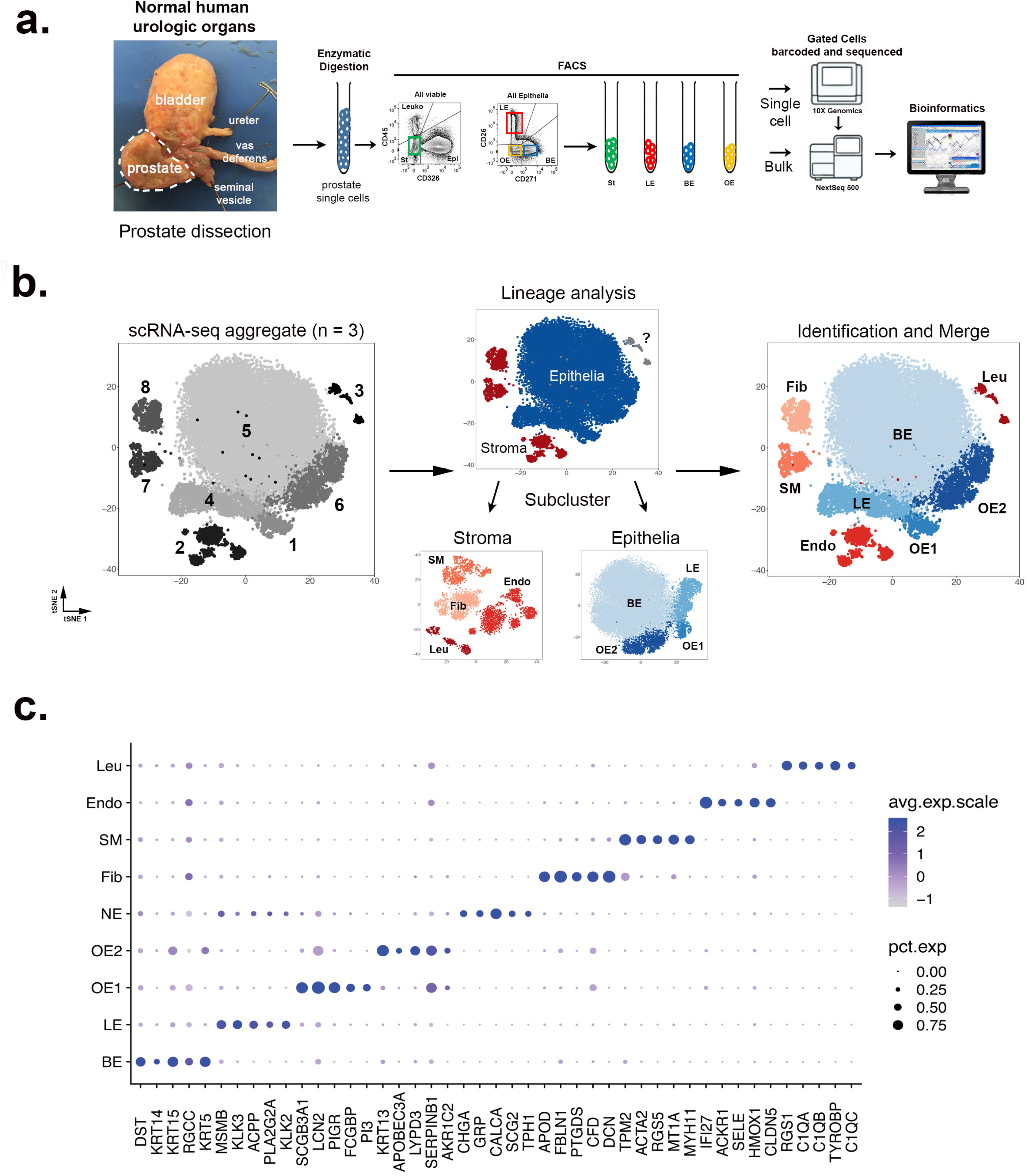
Identification of human prostate cell clusters with bulk and single cell RNA sequencing. (a) Schematic of human tissue collection and processing for bulk and single cell RNA sequencing. (b) Aggregated scRNA-seq data from three organ donor prostate specimens with sub-clustering into stroma, epithelia, and unknown lineages based on correlation with bulk sequencing data (Supplementary Figure 2). Clusters were identified and re-merged. (c) Dot plot of cluster-specific genes after *in silico* removal of stressed cells and supervised identification of neuroendocrine epithelia.

The stromal gate includes all cells negative for CD326 (pan-epithelia), CD45 (pan-leukocyte), and CD31 (endothelia). We bulk sequenced the triple negative stromal gate from each patient to generate epithelial-specific differentially expressed gene (DEG) sets for each epithelial population (Supplementary Figure 2b,c). Gene set enrichment analysis (GSEA) was performed to compare DEGs in our data set to other data sets of sorted prostate cells^7,8,12^. Luminal (CD26^+^) and basal (CD271^+^) epithelial transcriptomes from our study were highly concordant with those from other studies (Supplementary Figure 2d).

Familiar DEGs genes such as *KRT5*, *KRT14*, and *TP63* as well as novel genes such as *NOTCH4*, *LTBP2*, and *DKK1* characterize CD271^+^ basal epithelia. The CD26^+^ luminal epithelia are marked by familiar DEGs such as *KLK3*, *ACPP*, and *MSMB* as well as novel genes including *GP2*, *NEFH*, and *NPY*. Principal component analysis shows that the transcriptome of CD271^-^/CD26^-^ ‘other’ epithelia resembles that of CD271^+^ basal epithelia (Supplementary Figure 2a). However, several of the top ‘other’ epithelia DEGs include classic neuroendocrine lineage markers such as *CHGA* and *CHGB* and novel markers such as *LY6D*, *SCGB3A1*, and *PSCA*. Twenty significant DEGs in the three epithelial and one stromal gate are shown in Supplementary Figure 2e. Supplementary Table 1 includes the full list of cell type-specific DEGs.

Many of the significant DEGs in the ‘other’ epithelial gate are putative neuroendocrine cell markers, but the high frequency of cells in this gate argues against a pure population of neuroendocrine cells given their low frequency *in situ*. Before we could identify whether each FACS gate contained a heterogeneous population of cells, we employed single cell RNA sequencing (scRNA-seq) to establish an objective identity of each cell type.

### Single cell sequencing of the normal human prostate reveals unbiased cellular identities

Prostate specimens from 3 young male organ donors aged 18-31 were collected fresh, dissected, and digested into single cells. Single cell suspensions were then stained and sorted for viability by flow cytometry. Approximately 34,000 viable cells from each of the 3 specimens were loaded into a 10x Genomics Chromium controller for transcript barcoding (Figure 1a). After aggregating data from each specimen, 35,865 cells were barcoded with a normalized read depth of 22,729 per cell and an average of 1,356 genes detected per cell. Supplementary Table 2 provides the sequencing metrics for each sample.

We clustered the single cell transcriptomes with a modified version of the Seurat R pipeline^13^ (see Methods). The artifact of cellular stress created by dissociating solid tissues into single cell suspensions is an unavoidable issue that can be mitigated by the removal of affected cells prior to sequencing or *in silico*^14^. To identify and remove stressed cells from the analysis *in silico*, we built a bioinformatics tool based on a principle component analysis of an experimentally-derived stress signature, which detected stressed cells in nearly every cluster (Supplementary Figure 3). A prostate-specific stressed cell DEG list was subsequently derived and deployed to exclude stressed cells in future analyses (Supplementary Table 3). After ~10% of the total cells were removed due to a high stress signature, a tSNE plot of the remaining 28,606 non-stressed cells revealed eight clusters (Figure 1b). DEGs were generated for each cluster in Seurat for assigning identity.

To quantitatively assign the cellular identity of each cluster, we performed QuSAGE gene set enrichment analysis^15^. We first defined epithelial and stromal lineages by correlating the cluster transcriptomes to our bulk sequencing data (Supplementary Figure 4a). Once the broad lineages were identified (epithelia and stroma), each lineage was sub-clustered, and re-clustered for deeper identification. Prostate-specific fibroblast and smooth muscle transcriptomes have not previously been generated due to the inability to isolate these cell types. We therefore used four stromal cell gene ontology terms (muscle, fibroblast, endothelia, leukocyte) to characterize stromal sub-clusters (Supplementary Figure 4b). We then used bulk sorted prostate epithelial cell transcriptomes (CD26^+^ ‘luminal’, CD271^+^ ‘basal’, CD26^-^/CD271^-^ ‘other’) to identify epithelial sub-clusters (Supplementary Figure 4c). As shown in Figure 1b, clusters 2, 3, 7, and 8 were highly correlated with bulk sequenced stroma and identified as endothelia, fibroblasts, smooth muscle, and leukocytes, respectively (see Methods and Supplementary Figure 4b). Clusters 1, 4, 5, and 6 were highly correlated with bulk-sequenced epithelia. Cluster 5 displayed the highest correlation with bulk sequenced basal epithelia (BE). Clusters 1 and 6 most resembled the double negative (CD26^-^/CD271^-^) ‘other epithelia’ gate and were tentatively assigned labels ‘OE1’, and ‘OE2’. Cluster 4 displayed the highest correlation with bulk sequenced luminal epithelia (LE) (Supplementary Figure 4c). Figure 1c displays a dot plot of the top five cluster-enriched DEGs to highlight specificity for each cluster. A full list of genes filtered for cell type-specific expression for each cluster is shown in Supplementary Table 3.

Neuroendocrine cells are a rare prostate epithelial type *in situ*^6^. We did not identify a unique neuroendocrine cell cluster in the 24,450 sequenced prostate epithelial cells even when increasing the read depth to 75,000 reads/cell (the effect of read depth on cluster identification can be seen in Supplementary Figure 5). Because a complete normal neuroendocrine cell transcriptome has yet to be generated, we relied on a small number of putative marker genes to identify NE cells by a principle component-based approach (see Methods). We then developed a neuroendocrine cell score, identified 25 putative neuroendocrine cells, and created a detailed DEG list for future analyses (Supplementary Figure 6a-c and Supplementary Table 3). A novel marker gene for neuroendocrine cells that was discovered in our data set (SCG2) was tested in combination with a known neuroendocrine cell marker (CHGA) and confirmed *in situ* (Supplementary Figure 6d). Interestingly, several NE cells were labeled with either SCG2 or CHGA suggesting potential NE cell heterogeneity. A recent study demonstrated that NE cells are derived from a luminal progenitor^9^, which may explain why NE cells are mainly found within the luminal epithelial cluster (Supplementary Figure 6b).

### Single cell sequencing data improves FACS of human prostate cell types

With each cell type now identified objectively by its transcriptome, we next turned our attention to whether our current approach to isolating each of these cell types could be improved. The capture of multiple cell types in an individual flow cytometry gate diminishes the interpretation of outcomes in *ex vivo* experiments on cell type-specific function. To calculate the purity of our traditional flow cytometry gates (Figure 2a)^11^, we uniquely barcoded cells from each FACS gate using a single specimen and then aggregated the data. Figure 2b demonstrates the cell types present in each FACS gate, which is quantitated in Figure 2c. The fibromuscular stroma (FMSt) gate was 72% fibroblasts and smooth muscle with 28% endothelial contamination. The basal epithelia (BE) gate was largely homogenous, consisting of 93% BE cells. The luminal epithelia (LE) gate was highly contaminated with OE1 cells and the OE gate contained 50% BE cells, 24% OE1 cells, and 25% OE2 cells.

**Figure 2.**
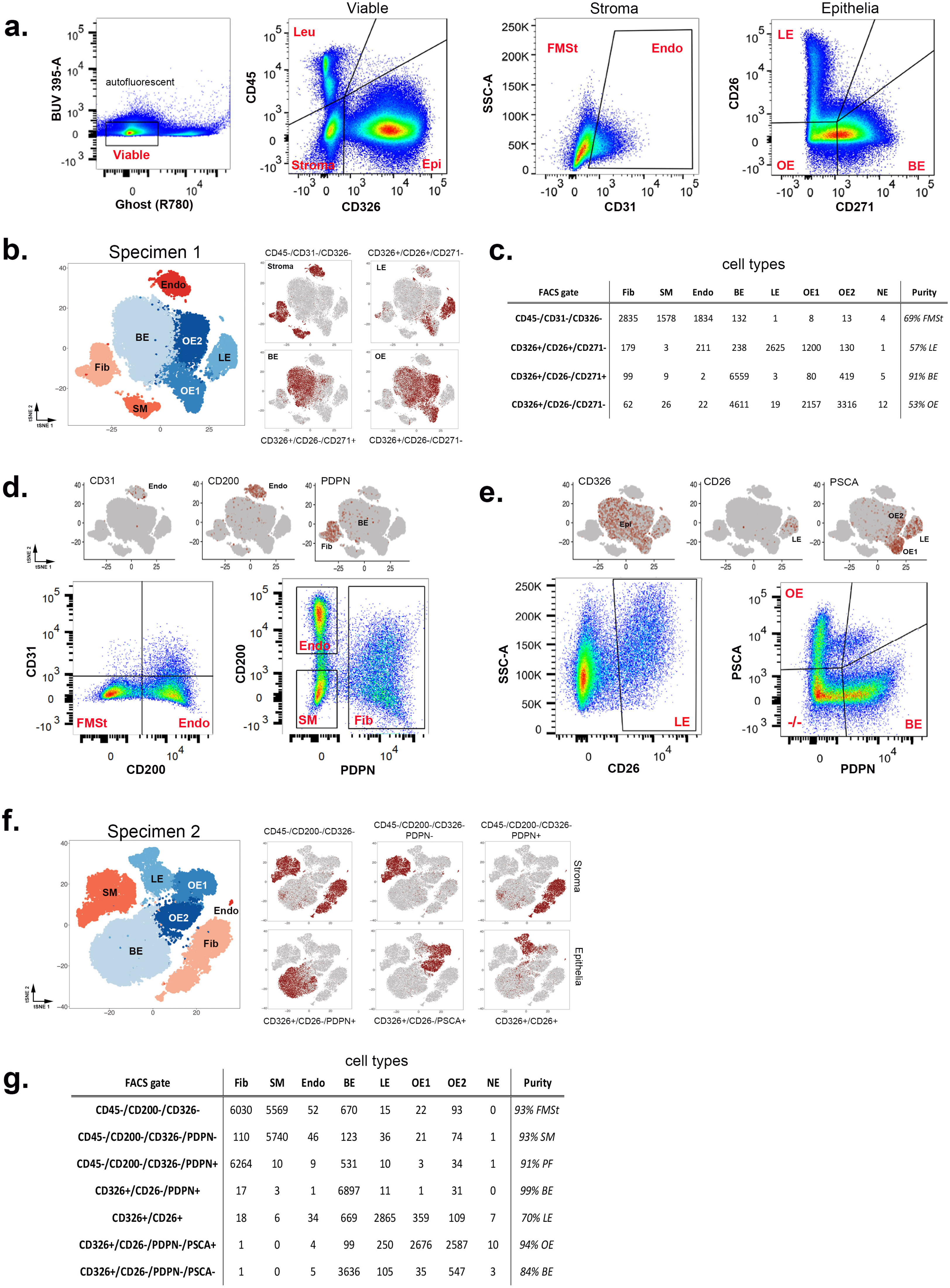
Optimization of flow cytometry for purification of stromal and epithelial subtypes. (a) Standard flow cytometry strategy for purification of prostate stroma and epithelial subtypes. (b) Barcoding of cells from traditional FACS gates shows breakdown of cell types within each gate. (c) Quantification of cells within barcoded FACS gates. (d) (left) CD200 labels 93% of endothelia that CD31 labels; (right) PDPN and CD200 separate endothelia (CD200^+^), fibroblasts (PDPN^+^) and smooth muscle (PDPN^-^). (e) PSCA was identified as a potential cell surface marker capable of isolating ‘other’ epithelial cells after CD26^+^ luminal epithelia are removed. (f, g) scRNA-seq of modified FACS gates on a new organ donor prostate specimen is used to demonstrate the increased purity of isolated stromal and epithelial cell types compared to traditional gates in panel a-c.

We increased the purity of FACS gated cells by using the scRNA-seq data set to identify novel cell surface markers. The primary contamination in the fibromuscular stroma is endothelia, leading us to conclude that CD31 is an inefficient endothelial marker in human prostate, even though it is widely used as such in mouse and human studies. To find a suitable replacement, we searched the endothelial cell cluster DEGs for a cell surface marker more inclusive than CD31 (PECAM1). CD200 is expressed in the majority of clustered endothelial cells (Figure 2d) and an antibody with multiple conjugation options is commercially available. Co-staining of human prostate single cells with CD31 (BV421) and CD200 (PE) reveals that 91% of CD31^+^ cells co-label with CD200 (Figure 2d), which led us to replace CD31 with CD200. We also searched the scRNA-seq dataset for a cell surface marker capable of separating fibroblasts from smooth muscle, which has not been feasible. After testing multiple options, Podoplanin (PDPN) was found to robustly label fibroblasts. Dual labeling of the CD45^-^/CD326^-^stromal gate with CD200 and PDPN shows three distinct gates on flow cytometry: CD200^+^ endothelia, PDPN^+^ fibroblasts, and CD200^-^/PDPN^-^smooth muscle (Figure 2d).

We next optimized a strategy for isolating pure epithelial cell types. The two most impure epithelial FACS gates are CD26+ luminal epithelia (contaminated with OE1), and the double negative (CD26^-^/CD271^-^) ‘other’ epithelia gate (contaminated with BE). Finding a positive marker for ‘other’ epithelia solves both issues. Accordingly, we mined scRNA-seq data to identify a novel cell surface antigen to separate basal, luminal, and other epithelia by FACS. After testing several options, PSCA was identified in the cluster-specific DEGs as a potential cell surface marker of both OE cell types (as well as a subpopulation of luminal epithelia) (Figure 2e). Flow cytometry with a PSCA antibody indicated that indeed ~50% of the CD26^+^ luminal epithelia were PSCA^+^ (data not shown), which necessitated the gating of PSCA^+^/CD26^+^ LE before gating of PSCA^+^ OE. PDPN was used to mark BE, which we previously showed largely overlaps with CD271^11^. As shown in Figure 2e, using PDPN for FACS gating of BE instead of CD271 is based on the fact that it can also be used to isolate the two stromal cell clusters (Figure 2d), thereby reducing the compensation issues associated with larger antibody panels.

To determine whether this newly optimized FACS scheme was superior to the traditional approach for purifying stromal (CD200^+^ endothelia and CD200^-^FMSt) and epithelial (PDPN^+^ BE, CD26^+^ LE, and PSCA^+^ OE) cells, we performed single cell sequencing on each new gate using a new young human prostate specimen (Figure 2f). First, we demonstrate increased purity of the FMSt population from 71% to 93% by reducing endothelial cell contamination (Figure 2f, g). Next, we improved purity of LE cells from 60% to 70% by reducing OE1 contamination. Purity of the BE gate (CD326^+^/CD26^-^/PDPN^+^) was improved from 93% to 99% and purity of the new OE gate (CD326^+^/CD26^-^/PDPN^-^/PSCA^+^) was 93% for OE1 and OE2 cells compared to 49% in the original CD326^+^/CD26^-^/CD271^-^ gate (Figure 2c). The triple negative (CD26^-^/PSCA^-^/PDPN^-^) CD326^+^ epithelial gate contained 80% BE and 16% OE2 cells not captured by PDPN and PSCA (Figure 2g). In summary, this improved antibody panel (CD45/CD326/CD200/CD26/PDPN/PSCA) for FACS gating will be instrumental in the functional characterization of purified human prostate epithelial and stromal cell types.

### Identification and isolation of prostate stromal cell subtypes

We next focused on classifying stromal cell type identities by sub-clustering 4,156 stromal cells (endothelia, fibromuscular stroma, leukocytes) from three young organ donor prostate specimens (Figure 3a). DEGs were generated for stromal sub-clusters (Figure 3b). The putative smooth muscle cluster expressed high levels of actin (ACTA2) and myosin (MYH11, MYL9, TPM2) genes while the putative fibroblast cell type expressed high levels of paracrine signaling factors such as growth factors (FGF2, FGF7), prostaglandins (PTGDS, PTGS2), and WNT pathway regulators (RSPO3,SFRP2). We used FACS to isolate PDPN^-^ and PDPN^+^ stroma as well as CD200^+^ endothelia from 3 separate young organ donor prostate specimens and performed qPCR on known and novel cluster-specific DEGs to confirm the scRNA-seq data. The results demonstrate selective expression for each DEG in smooth muscle, fibroblasts, and endothelia (Figure 3c). Leukocytes were too low in these normal specimens to sort out sufficient numbers for qPCR and were therefore excluded from the comparison. To gain a better understanding of prostate cellular function, stromal sub-cluster transcriptomes were used to run QuSAGE against C2 curated genesets from MSigDB^16,17^ (Supplementary Table 4). Ten top pathways of the KEGG subset^18^ of C2 pathways are displayed for smooth muscle and fibroblast cell types in Figure 3d. Of note, fibroblasts show a high enrichment for ‘protein export’ suggesting a putative paracrine function and smooth muscle show enrichment of contraction and metabolism pathways. Finally, we tested antibodies for immunohistochemical detection of each cell type *in situ* and found that Myosin 11 (MYH11) and Decorin (DCN) are enriched in smooth muscle and fibroblast cell types, respectively, in human prostate tissue (Figure 3e).

**Figure 3.**
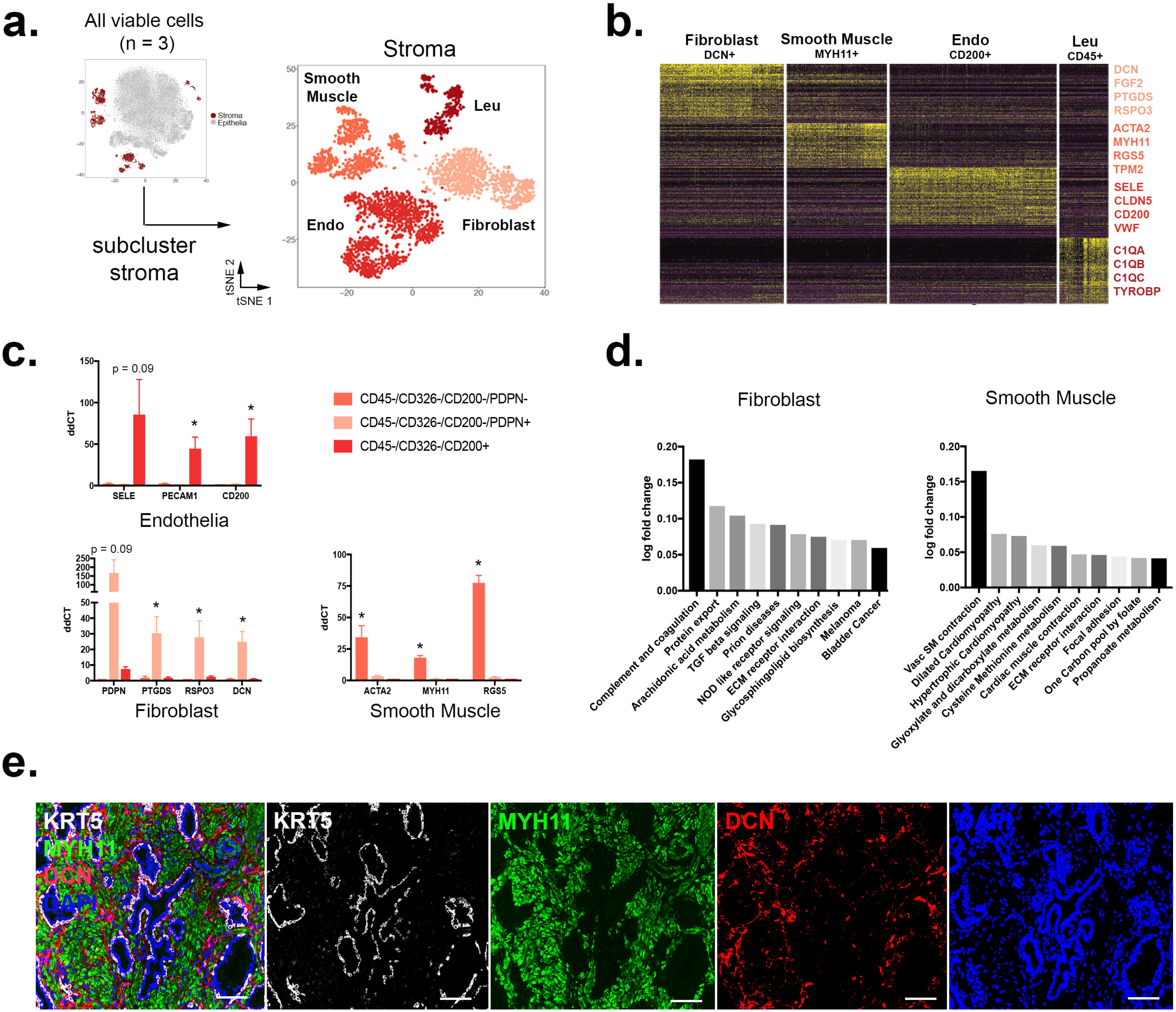
Identification and isolation of pure stromal subtypes in the normal human prostate. (a) Single cell RNA sequencing data aggregated from three normal prostate specimens sub-clustered into the stromal lineage. (b) Heatmap of top 100 differentially expressed genes in each stromal subcluster with highlighted DEGs suggesting putative identities. (c) qPCR of FACS-isolated stromal subtypes from three organ donor prostate specimens demonstrates the enrichment of cell type-specific DEGs. (d) GSEA of non-endothelial stromal populations compared to KEGG pathways. (e) Immunofluorescent labeling of smooth muscle (MYH11), fibroblasts (DCN), and basal epithelia (KRT5). * = p ≤ 0.05; Scale bar = 100μm.

### Identification and isolation of epithelial cell subtypes

We next objectively defined epithelial cell type identities by sub-clustering the 24,450 epithelial cells from three organ donor specimens. DEGs of each cluster confirm four epithelial cell types (Luminal KLK3^+^, Basal KRT14^+^, OE1 SCGB1A1^+^, and OE2 KRT13^+^) (Figure 4a, b). Viable neuroendocrine epithelia were too infrequent to cluster independently and were detected through principle component analysis (see Methods and Supplementary Figure 5). We confirmed the gating scheme for isolating CD26^+^ luminal, PDPN^+^ basal, and PSCA^+^ ‘other’ epithelia shown in Figure 2e by performing qPCR on known and novel cluster-specific DEGs (Figure 4c).

**Figure 4.**
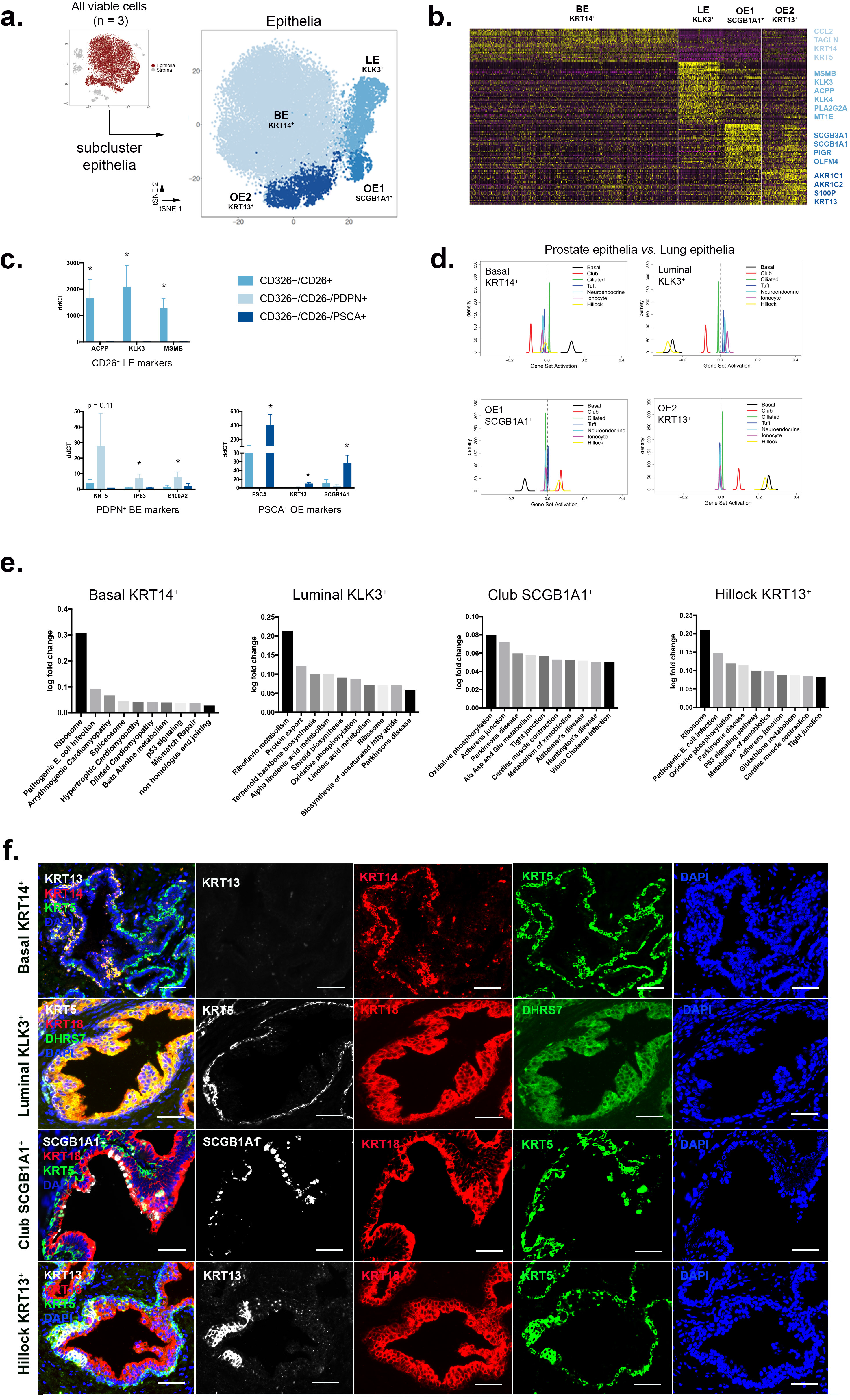
Identification and isolation of pure epithelial subtypes in the normal human prostate. (a) Single cell RNA sequencing data aggregated from three normal human prostate specimens sub-clustered into the epithelial lineage. (b) Heatmap of top 100 differentially expressed genes in each epithelial subcluster with highlighted DEGs. (c) qPCR of FACS-isolated epithelial subtypes from three organ donor prostate specimens demonstrates the enrichment of cell type-specific DEGs. (d) GSEA of four human prostate epithelial cell types compared to mouse lung epithelial cell types. (e) KEGG pathways enriched in epithelial cell types. (f) Immunofluorescent labeling of basal epithelia (KRT5), luminal epithelia (DHRS7), club epithelia (SCGB1A1) and hillock epithelia (KRT13). * = p ≤ 0.05; Scale bar = 50μm.

SCGB proteins, or ‘secretoglobins’, are highly expressed by respiratory tract club cells^19^ but have also been detected in human prostate^20^. To determine whether SCGB1A1^+^ prostate epithelia are transcriptionally similar to lung club cells, we performed QuSAGE with our human prostate epithelial scRNA-seq data compared to a scRNA-seq dataset from mouse lung epithelia^21^ (Figure 4d). These data demonstrate a strong correlation between SCGB1A1^+^ prostate epithelia and lung Scgb1a1^+^ club cells as well as lung Krt13^+^ hillock cells. KRT5^+^/KRT13^+^ prostate epithelia display a strong positive correlation with lung Krt5^+^ basal and Krt13^+^ hillock cells. KRT5^+^/KRT14^+^ prostate basal epithelia are highly correlated with lung Krt5^+^ basal cells. KLK3^+^ prostate luminal epithelia are not strongly correlated with any of the mouse lung cell types, but were highly correlated with lung AT2 secretory cells in a human scRNA-seq lung data set^22^ (data not shown).

To better understand potential functions for each cell type, epithelial subcluster transcriptomes were used to run QuSAGE against C2 curated genesets from MSigDB^16,17^ (Supplementary Table 4). Ten top pathways of the KEGG subset^18^ of C2 pathways are displayed for each cell type in Figure 4e. KEGG pathways strongly correlating with Luminal KLK3^+^ epithelia include lipid and steroid metabolism while KRT14^+^ basal epithelia display a variety of enriched pathways including proteasome, ribosome, and amino acid metabolism. SCGB1A1^+^ and KRT13^+^ ‘other’ epithelial cell types both displayed a strong correlation with immunomodulatory pathways. A full list of epithelial cell type-specific DEGs is provided in Supplementary Table 3.

As shown in Figure 4f, we optimized immunohistochemical detection protocols each cell type *in situ* and found that KRT14^+^ basal epithelia are KRT13^-^ and represent a subpopulation of KRT5^+^ basal epithelia. DHRS7 was a novel gene detected in KLK3^+^ luminal epithelia and marks a portion of KRT18^+^ luminal epithelia. SCGB1A1 positively marks a small population of KRT5-luminal-like prostate epithelia. Finally, KRT13^+^ epithelia co-express KRT5, confirming GSEA analysis showing a positive correlation with basal cells of the lung (Figure 4d). These data establish distinct KRT5^+^ basal epithelial cell types that are discriminated by co-staining with KRT14 or KRT13.

### Novel epithelial cell types are enriched in the urethra and peri-urethral prostate zones

Benign and malignant prostate diseases are largely restricted to the transition/central and peripheral zones, respectively^23^. A deeper understanding of each zone’s cellular anatomy may shed light on why these diseases predominate in distinct regions. After characterizing the molecular identity of novel human prostate stromal and epithelial cell types, we examined whether they are differently distributed across anatomical zones. Accordingly, we dissected the transition and central zones from the peripheral zone as shown in Figure 5a. For two specimens, each anatomical zone was digested into single cells, sorted for viability, and processed for scRNA-seq. To determine the natural incidence of each cell type in each anatomical zone, we superimposed the cells in each zone onto the aggregated data. Quantification of the results revealed that the transition/central zones are enriched for club-like SCGB1A1^+^ (OE1) and KRT13^+^ (OE2) epithelia but luminal epithelia are low (Figure 5b). To confirm these data, we performed flow cytometry with our optimized antibody panel to quantitate the number of PSCA^+^ other epithelia in TZ/CZ *vs*. PZ from 5 more young organ donor specimens (Table 1 and Figure 5c). Quantification of the FACS data confirmed that PSCA^+^ ‘other’ epithelia are enriched as a percentage of epithelia in the transition/central zones while CD26^+^ luminal epithelia are enriched as a percentage of epithelia in the peripheral zone (Figure 5d).

**Figure 5.**
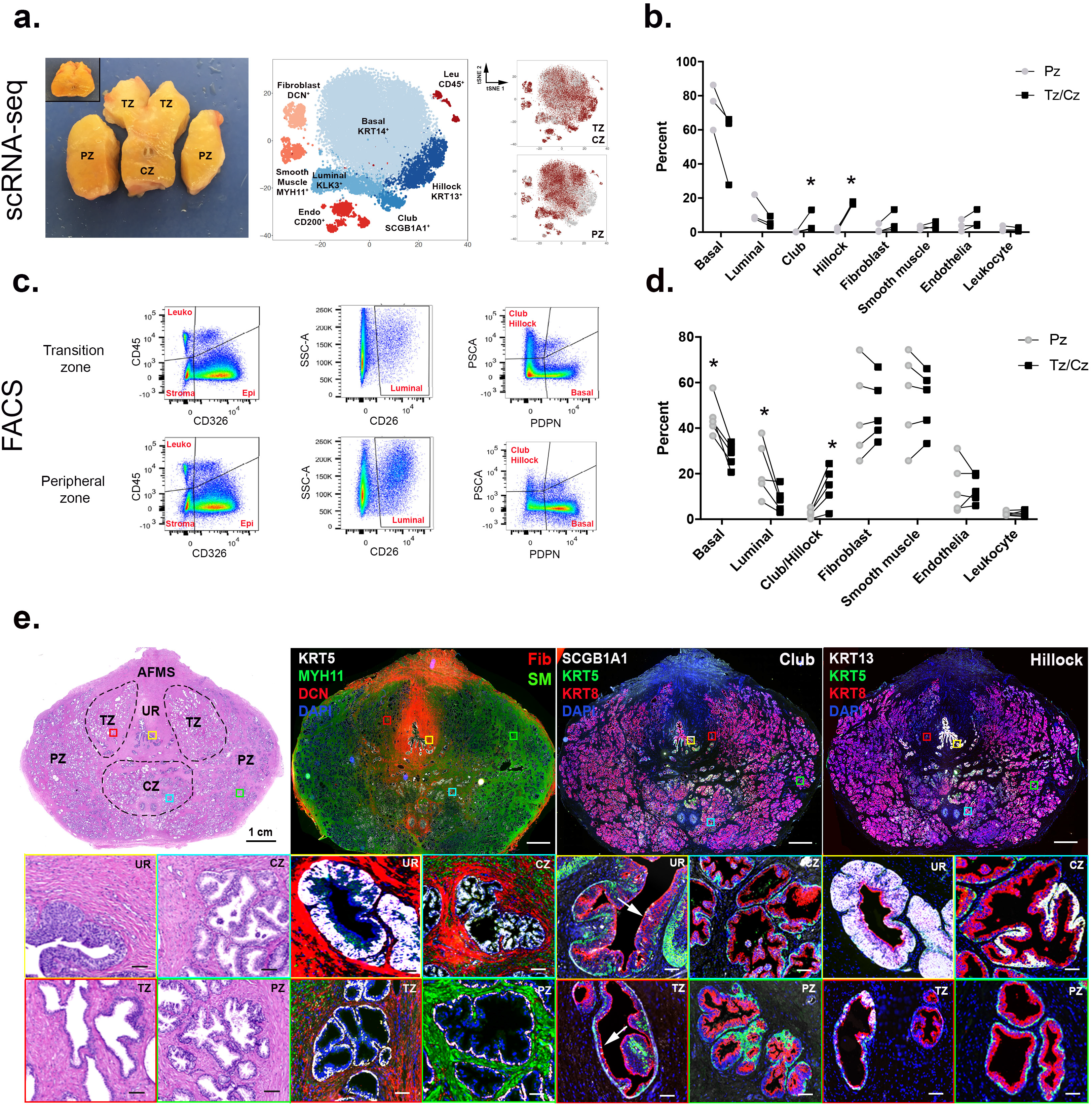
Anatomical location of epithelial and stromal cell types in the normal human prostate. (a) The transition and central zones of the prostate were dissected away from the peripheral zone from three young organ donors for scRNA-seq (pre-dissected tissue inset). (b) Quantification of scRNA-seq-identified cell types after segregation by anatomical zone from 3 patient aggregated data. (c) Representative FACS analysis of epithelia from transition and peripheral zone tissue from five young organ donors after segregation by anatomical zone. (d) Quantification of FACS data on zonal enrichment of cell types. (e) Immunofluorescence of prostate whole mount sections displays enrichment of novel epithelial cell types in the central and transition zones and the urethra and a concentration of fibroblasts in the peri-urethral and central zone regions. * = p ≤ 0.05; Scale bar = 100μm.

**Table 1.**
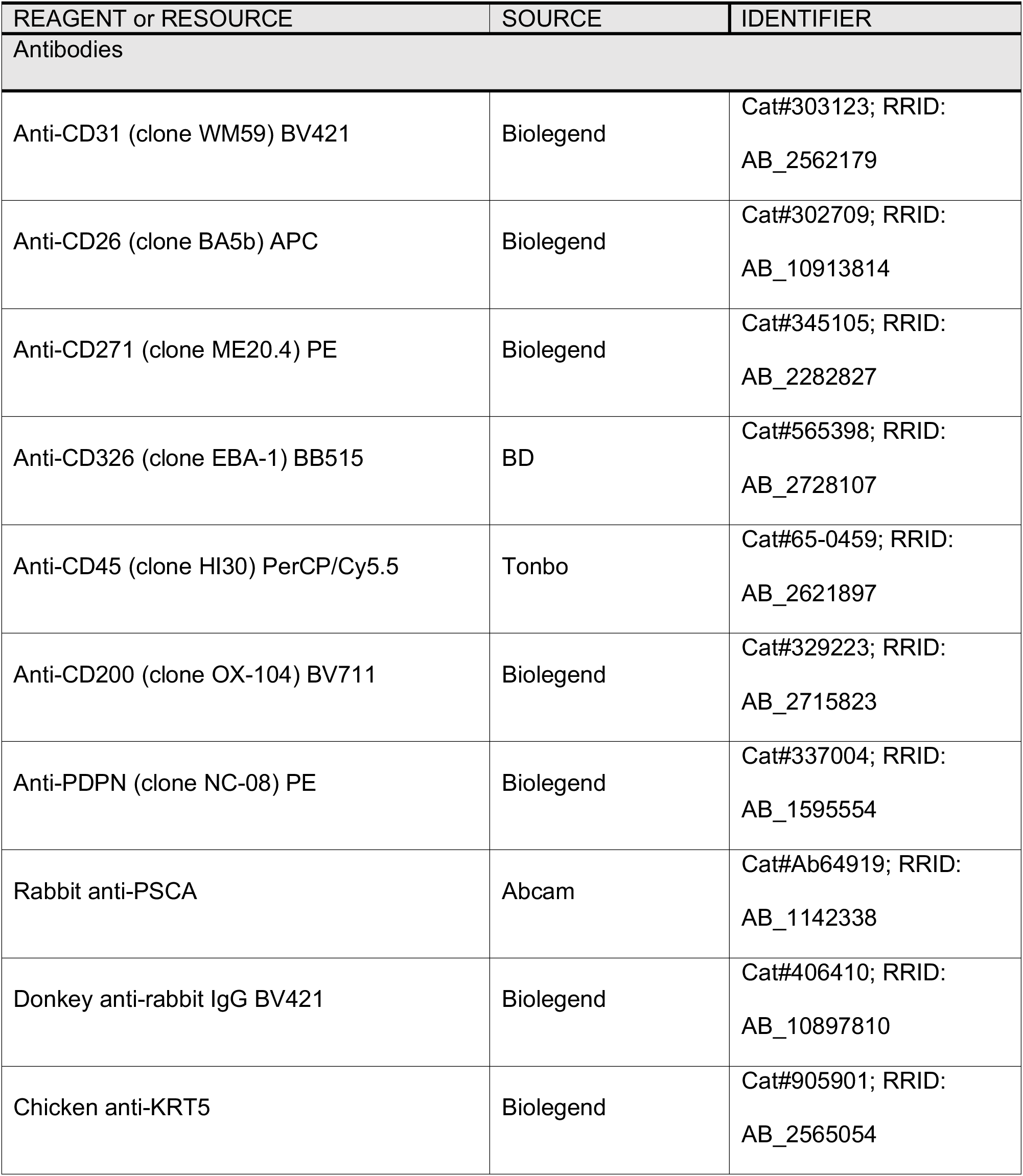
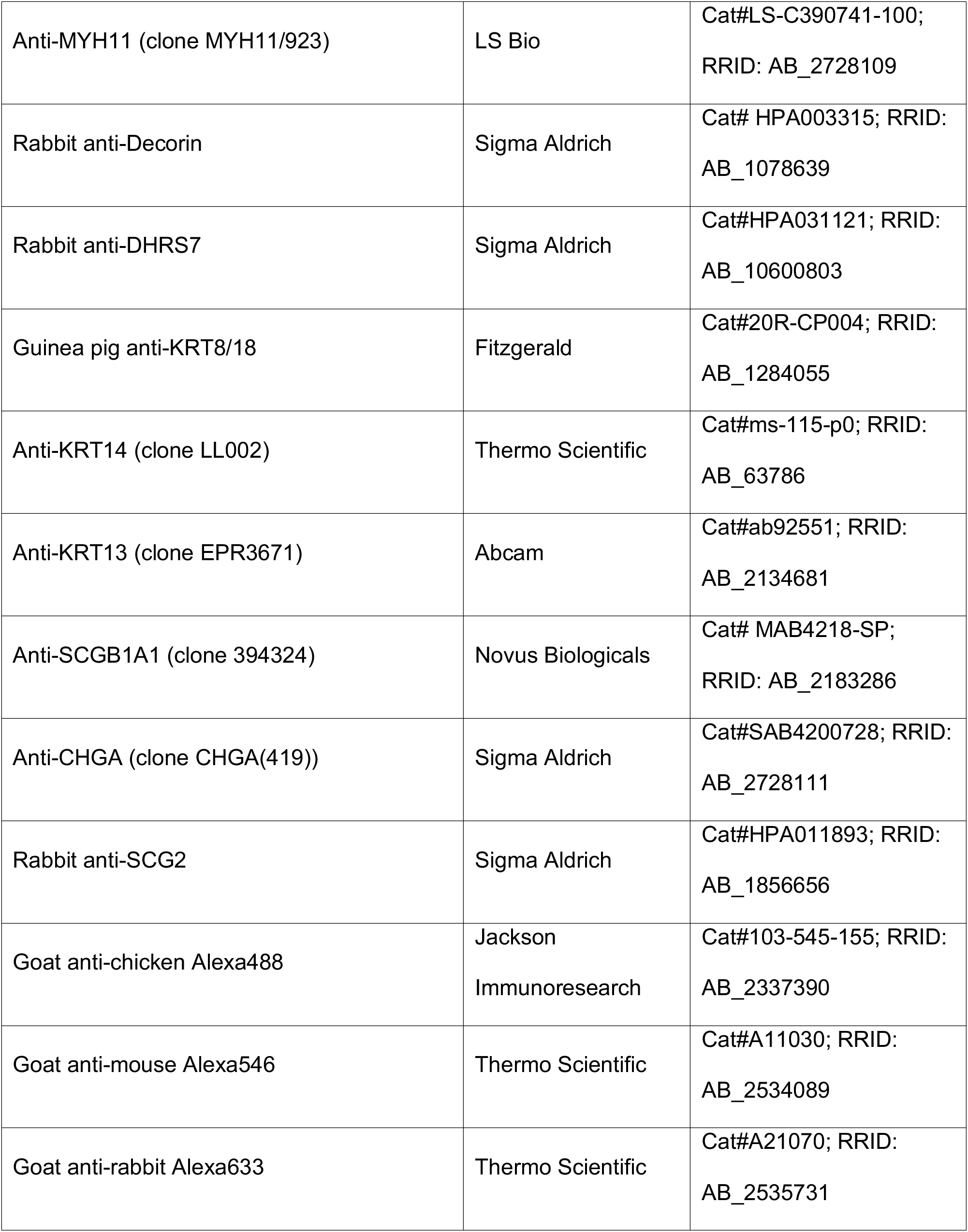
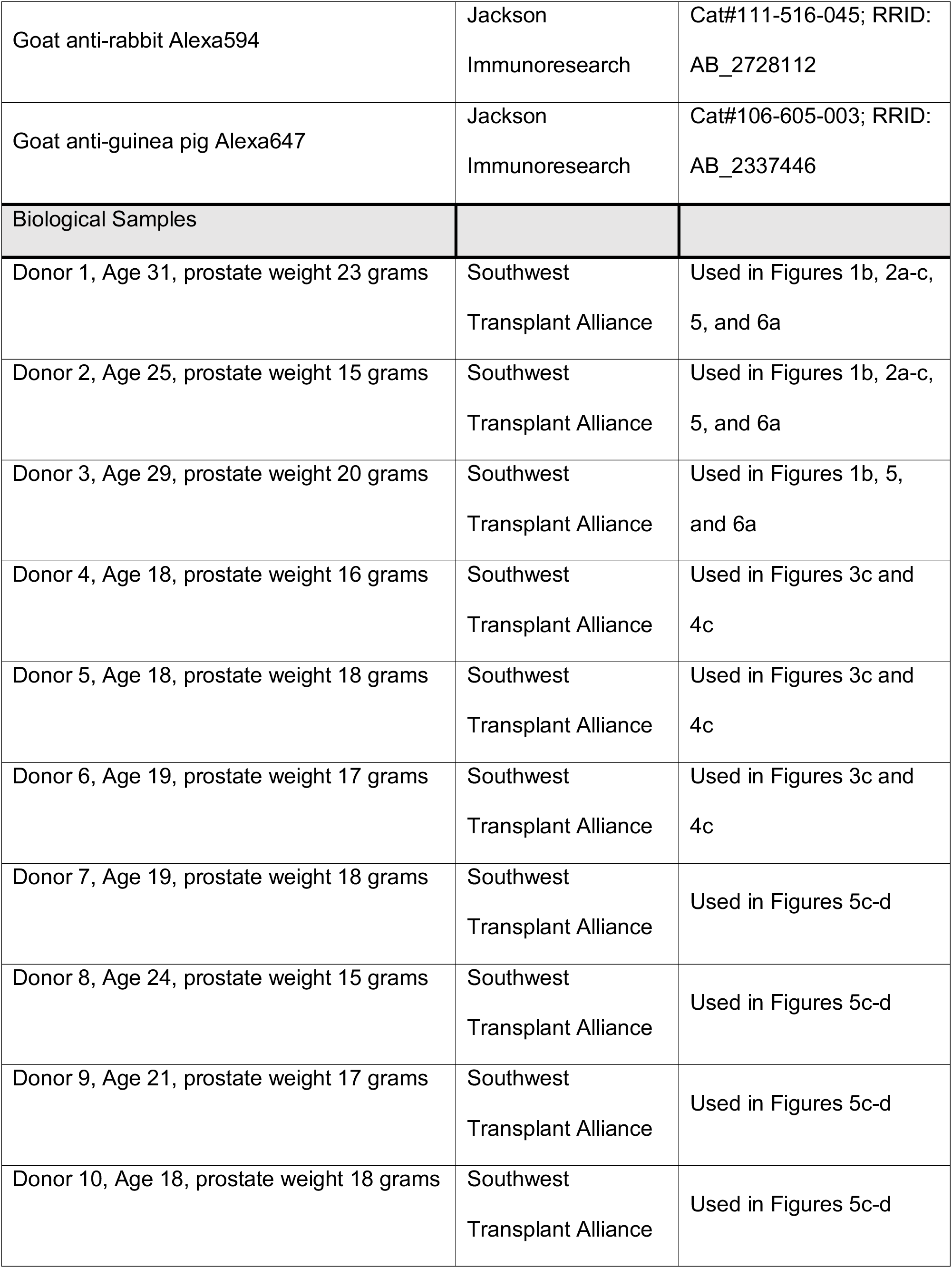
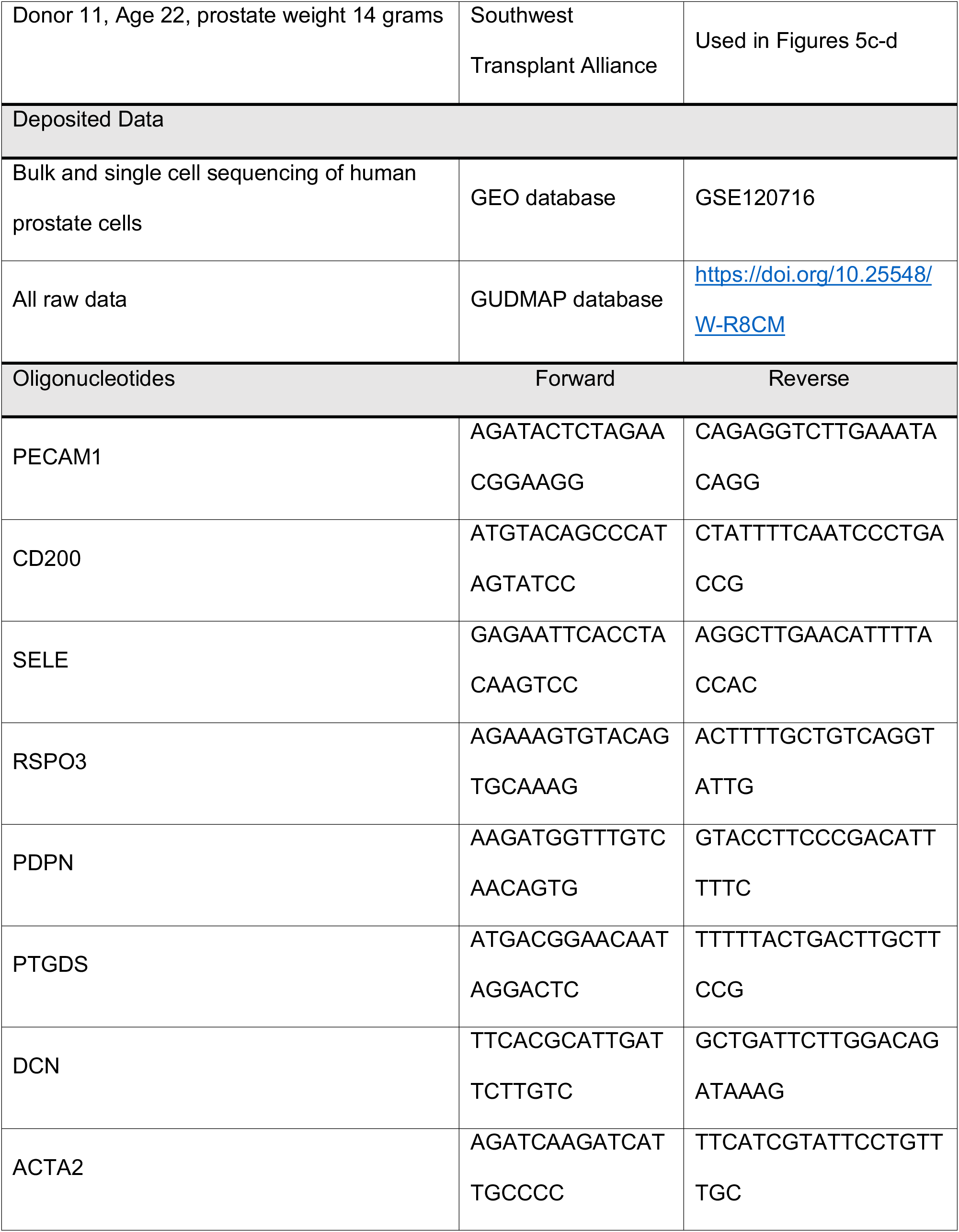
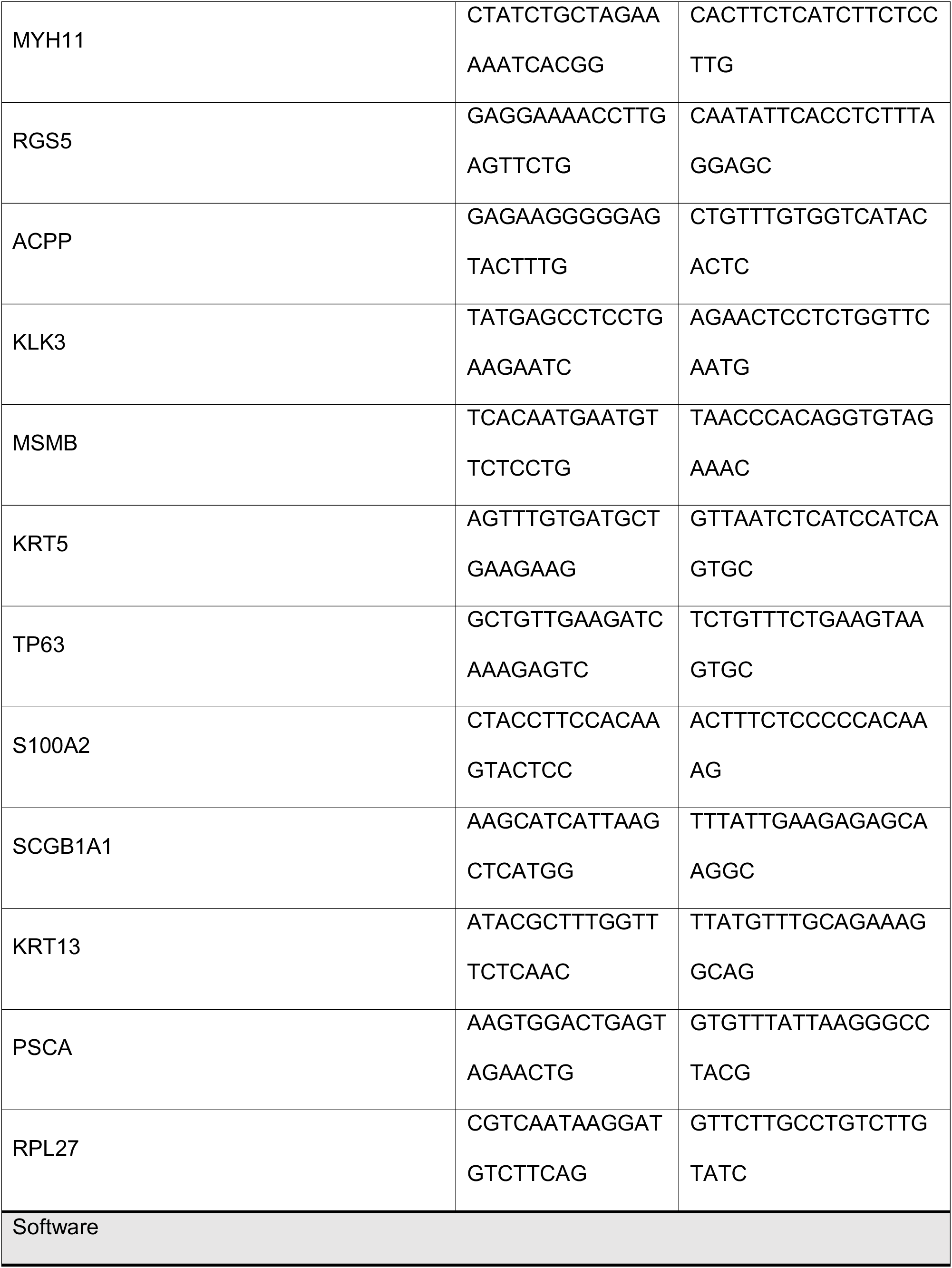
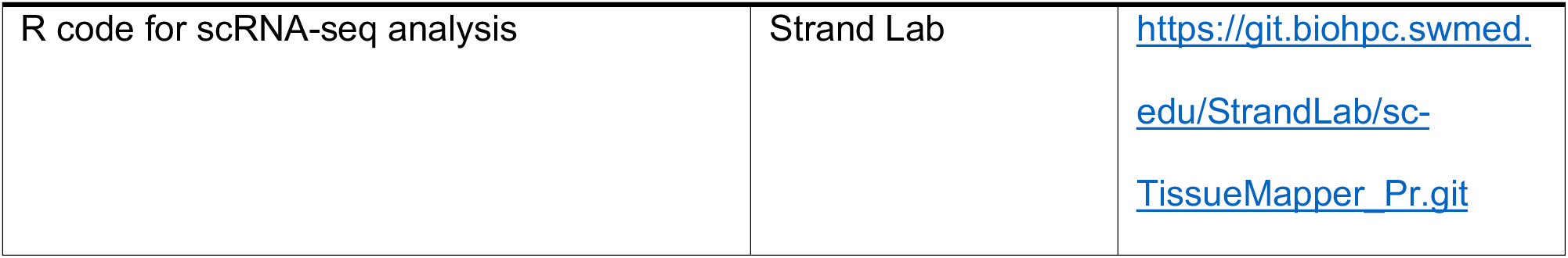
Key resources for entire study.

To confirm these trends *in situ*, we performed triple immunofluorescence with markers of each cell type on whole transverse sections of the normal human prostate by collecting tiled images and stitching them together. KRT5^+^/KRT14^-^/KRT13^+^ epithelial cells are abundant in the prostatic urethra and collecting ducts as well as the central zone surrounding the ejaculatory ducts, but are rare in the peripheral zone (Figure 5e). KRT5^-^ /KRT8^-^/SCGB1A1^+^ cells are abundant in the prostatic urethra and collecting ducts and rare in the prostate. Fibroblasts are common in the pre-prostatic region surrounding the urethra, anterior fibromuscular stroma (AFMS), and the transition and central zones. Smooth muscle myocytes are the predominant peripheral zone stromal cell type.

The anatomical distribution of particular stromal and epithelial cell types in the proximal (transition/central zones) and distal (peripheral) prostate could underlie the regional incidence of benign and malignant diseases. Because both diseases have been suggested to arise from putative stem cells, the lineage hierarchy of the mouse and human prostate has been studied extensively. Multipotent progenitor cells of the primitive urethra and proximal prostatic ducts give rise to prostate buds early in development^24^ and prostate glands in the adult^25^, respectively, but their identity is incompletely understood. We performed pseudotime analysis^26^ to build single cell trajectories to gain a better understanding of the dynamical relationships among club, hillock, basal, and luminal epithelial cell types. Figure 6a demonstrates a diversion of luminal and club/hillock cell types from basal cells, which may be analogous to the hierarchy of the lung where a multipotent Krt5^+^ basal epithelial cell gives rise to all differentiated cell types while club and hillock cells are more restricted progenitors^21^. An illustrated atlas of the regional distribution of objectively defined cell types in the normal human prostate and prostatic urethra is displayed in Figure 6b.

**Figure 6.**
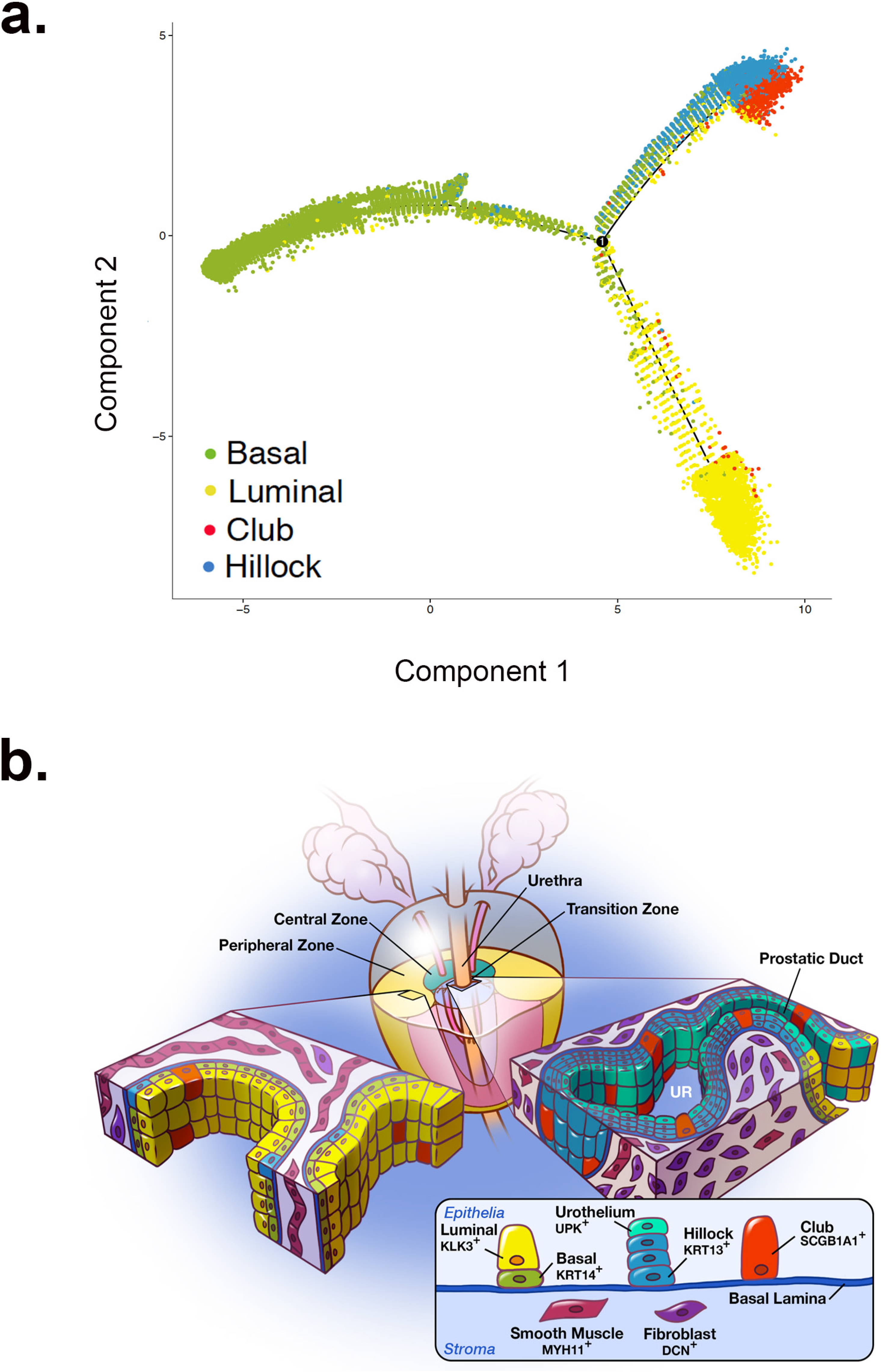
Cellular anatomy and hierarchy of the normal human prostate. (a) Pseudotime analysis of human prostate epithelia. (b) Illustrated cellular atlas of human prostate zonal anatomy displays the enrichment of hillock KRT13^+^ and club SCGB1A1^+^ epithelia in the urethra and proximal prostate ducts.

## Discussion

The cellular origins of BPH^27^ and prostate cancer^28^ are still unknown. To understand how cellular composition and cell type-specific gene expression change in disease, a proper control must be set. Routine access to normal adult human specimens is limited so alternative controls such as ‘normal adjacent’ areas of diseased specimens have been substituted without regard for field defect^29^. We present here an objective characterization of the molecular identity and location of each cell type in the normal adult human prostate, as well as a validated experimental tool to isolate pure cell types.

Cell identity has been historically based on a small set of marker genes. In fact, most lineage tracing- or flow cytometry-based studies rely on a single marker to define cell type. Single cell RNA sequencing (scRNA-seq) has revolutionized the idea of cell identity by providing an unbiased genetic signature across a set of cells^30^. Using scRNA-seq, we derived a molecular identity of 5 epithelial cell types and 2 stromal cell types in young adult human prostate (Figure 1). These data were then used to find optimal cell surface markers for enriching defined cell types by FACS, and also to develop immunostaining protocols for genes that uniquely identify each cell type *in situ* (Figures 2–5). The results confirmed the existence of previously described prostate stromal cell types (fibroblast and smooth muscle) and previously described prostate epithelial cell types (basal, luminal, neuroendocrine). However, we expanded our understanding of the identity, spatial location, and putative function of each cell type by providing a comprehensive transcriptomic signature. These data also led to the discovery of two previously unrecognized epithelial cell types marked by high expression of SCGB1A1 and KRT13 and an anatomical enrichment in the prostatic urethra and proximal prostatic ducts.

Prostate SCGB1A1^+^ cells are similar in morphology and transcriptomic profile to Clara, or ‘club’, cells (Figure 4d)^21^, which account for ~20% of the epithelial lining of the respiratory tract and are concentrated in the tracheal trunk^22^. Club cells are a non-ciliated, non-mucous, cuboidal secretory cell type that express anti-microbial, anti-viral, and anti-inflammatory proteins^31^. Although SCGB1A1 was previously shown to be expressed in the human prostate when examining whole tissue extracts^20^, it was not known to be a marker of a unique cell type. Prostate club cells are similar to lung club cells in their enrichment of immunomodulatory programs (Figure 4, Supplementary Tables 4 and 5), but their function in the prostate or prostatic urethra has not been tested.

Prostate KRT13^+^ cells are similar in morphology and transcriptomic profile to ‘hillock’ basal cells of the lung^21^. Prostate hillock cells are also concentrated in the prostatic urethra and proximal prostatic ducts (Figure 5). Cells with KRT13 expression were previously shown to be rare in the adult prostate, but abundant in fetal prostate, although it is unclear whether hillock cells populate the fetal urogenital sinus. KRT13^+^ cells are enriched in localized prostate tumors and in stem-like cells that display androgen resistance and a capacity for branching morphogenesis^32^. Intriguingly, the top genes associated with the KRT13^+^ cell type are members of the androgen metabolism pathway (AKR1C1 and AKR1C2), which have been implicated in the development of castrate resistant prostate cancer^33^. Until now, it has been assumed that KRT13 expression in prostate disease was simply increased in basal or luminal cell types. These new data suggest an intriguing hypothesis that a hillock cell type may be enriched in tumors.

Lung club and hillock cells can also act as progenitors for differentiated cell types^19,21,31^. Prostate club and hillock cells express high levels of PSCA (Figure 2). PSCA^+^ cells are enriched in prostate cancer^34^, but their full identity has not been firmly established. This is important because half of prostate luminal epithelia also express PSCA (Figure 2e). PSCA^+^ epithelial progenitors are also enriched in the proximal (peri-urethral) prostate of the mouse^35,36^, but this had not been confirmed in humans due to notable anatomical differences between mouse and human prostate^37,38^. These anatomical similarities could be confirmed if functional analyses show that prostate PSCA^+^ club and hillock cell types display multipotency as is found in the proximal lung^21^.

Club and hillock epithelial cell types could also play a role in BPH. The prostate buds off the urethra during development and subsequently undergoes branching morphogenesis into a ‘ductal tree’^24,39,40^. The adult human prostate displays 25-30 independent ductal structures connect separately to the urethra^41^. Clonal mapping of the human prostate shows that 95% of the progenitors that produce proximal to distal clones are found in the main trunks of these juxta-urethral ducts^25^. Using laser capture microdissection of the juxta-urethral trunk *vs*. distal prostate glands, Moad *et al* proposed that bipotent basal epithelial progenitors are enriched at the proximal prostate-urethral junction and are largely responsible for the homeostasis of the adult prostate epithelium^25^. The comprehensive cellular atlas produced here shows that the urethra and proximal ductal trunks of the prostate are predominantly composed of club and hillock cells (Figure 5). The characterization of marker genes and cell surface antigens capable of identifying these cell types *in situ* and purifying them for *ex vivo* study should facilitate the determination of whether these cells can also act as progenitors in normal epithelial homeostasis or whether they can act as progenitors in a putative stem cell disease such as BPH^42^.

Human tissue research relies on the isolation of cell types with cell surface markers, but the purity of the gated cells has only been inferred from bulk transcriptomic analysis which can conceal impurities through averaged gene expression. By identifying cellular subpopulations within FACS gates with scRNA-seq, we demonstrate that the purity of traditional gating schemes could be improved (Figure 2).

The gating of basal epithelia with either CD271 or PDPN produces >93% purity. Our previous work shows that most of these basal markers overlap and can be used interchangeably^11^. The initial gating of luminal epithelia with CD26 yielded a surprisingly low purity at 60% due largely to contamination with SCGB1A1^+^ club cells (Figure 2a-c). This gate was likely drawn too strictly, failing to account for spreading after the addition of antibody. However, this observation led to the realization that the CD38^lo^ (likely the same as CD26^lo^) ‘luminal’ cells described by Liu *et al* are likely enriched with SCGB1A1^+^ club cells, supported by the high enrichment score seen when comparing the CD38^lo^ transcriptome with the CD26^-^/CD271^-^ ‘other’ epithelia bulk transcriptome (Supplementary Figure 2d)^8^. This raises the intriguing possibility that the expansion of CD38^lo^ cells near sites of inflammation may be an expansion of club cells, which display anti-inflammatory and regenerative activity in the damaged lung^31^. Traditionally, the epithelial cell phenotype that expands during inflammation or injury has been described as an ‘intermediate’ cell type enriched for expression of KRT19 and sharing expression of luminal (KRT18) and basal (KRT14) cell types^4,43^. However, various forms of stress such as luminal anoikis, inflammation, or obesity can drive multipotent basal progenitors to give rise to luminal epithelia in the adult^44–46^. Comparing the comprehensive dataset generated here to a murine single cell data set will facilitate the tracing of definitive lineages to determine whether the response to injury is an expansion of a particular cell type or a transition between cell state, or both.

In addition to dramatically improving the isolation of each epithelial cell type, we for the first time demonstrate the ability to isolate pure populations of prostate stromal cell types. The first step towards this achievement was the recognition that the stromal gate was severely contaminated with endothelia due to the poor performance of the CD31 cell surface antibody. We first noted the contamination in our bulk sequencing of stroma, which displayed a high number of endothelial genes in the top DEGs (Supplementary Figure 2). Our first scRNA-seq experiment confirmed a 28% endothelial cell contamination of the stroma and also revealed CD200 as a potentially superior marker. The replacement of CD31 with CD200 improved the purity of the stromal gate from 72% to 93% (Figure 2). scRNA-seq also revealed PDPN as a positive marker of fibroblasts, which was used again on new FACS gates to confirm a 91% purity of PDPN^+^ fibroblasts and a 93% purity of PDPN^-^ smooth muscle (Figure 3f,g).

Previous studies have suggested the existence of at least four stromal cell types in the mouse prostate based on morphology, anatomical position and the expression of individual markers, including a population of ‘interstitial fibroblasts’ marked by Gli1^47^. A deeper analysis of these human data may reveal further fibroblast and smooth muscle subtypes similar to these mouse studies. The most striking discovery in the stroma was that the paracrine factors long thought to regulate prostate organogenesis such as WNTs, FGFs, and prostaglandins are predominantly expressed by fibroblasts and not by smooth muscle^48^. The concentration of DCN^+^ fibroblasts in the peri-urethral transition zone and peri-ejaculatory duct central zone could implicate these cells in the pathogenesis of BPH and should be examined further (Figure 5e).

Consortiums such as the Human Cell Atlas (HCA) and the GenitoUrinary Development Molecular Anatomy Project (GUDMAP) are efforts to provide markers for the identification of cell types in order to understand their functional interaction in normal organs. These efforts are a necessary foundation for a deeper understanding of disease^49,50^. Our study provides the deepest understanding to date of cell types found in the normal human prostate and prostatic urethra as well as their anatomical positions. The tools to identify and localize of every cell type in the normal human prostate is a valuable resource that establishes a baseline for all future studies of prostate disease.

## Methods

### Human prostate collection

Prostate specimens used in this study were obtained from 11, 18-31 year old male organ donors whose families were consented at the Southwest Transplant Alliance from March 2017 to April 2018 under IRB STU 112014-033. After transplantable organs were harvested, a cystoprostatectomy was performed and the specimen was transported to UT Southwestern Medical Center for processing. The prostate was dissected away from the bladder, and further dissected into anatomical zones as represented in Supplementary Figure 1. The average age was 22 and the average prostate size was 17 grams. Details for each specimen and its usage in associated figures are shown in Table 1.

### Tissue processing

Fresh tissue samples less than 24 hours post-mortem were transported in ice-cold saline and immediately dissected into portions for 1) flash freezing in liquid nitrogen, 2) fixation in 10% formalin followed by paraffin embedding, and 3) a 4 hour enzymatic digestion into single cells at 37ºC using 5 mg/ml collagenase type I (Life Technologies), 10μM ROCK inhibitor Y-27632 (StemRD), 1nM DHT (Sigma), 1mg DNAse I, and 1% antibiotic/antimycotic solution (100X, Corning) in HBSS^51,52^. Single cells were filtered, resuspended, and incubated with antibodies for flow cytometry.

### Flow cytometry

Viable human prostate cells were isolated by fluorescence activated cell sorting (FACS) for bulk and single cell sequencing in the UT Southwestern CRI Flow Cytometry Core on a BD FACSAria FUSION SORP flow cytometer and analyzed with FlowJo software as previously published^11^. New antibody panels based on single cell data were built using titration and fluorescence minus one experiments. Table 1 displays information on antibodies used for flow cytometry.

### Immunohistochemistry

Fluorescent immunohistochemistry was performed as described previously^53^. In brief, 5 μm paraffin sections were deparaffinized in xylene and hydrated through a series of ethanol washes. Heat mediated antigen retrieval was performed by boiling slides in 10 mM sodium citrate (pH 6.0) for 20 min in a conventional microwave oven. Tissues were washed with a solution containing 25 mM Tris-HCl, pH 7.5, 140 mM NaCl, 2.7 mM KCl, and 0.1% Tween-20 (TBSTw) and non-specific binding sites were blocked for 1 hr in TBSTw containing 1% Blocking Reagent (Roche Diagnostics, Indianapolis, IN), 5% normal goat sera, and 1% bovine serum albumin fraction 5 (RGBTw). Tissues were incubated overnight at 4°C with primary antibodies diluted in RGBTw. Tissues were washed several times in TBSTw and incubated with secondary antibodies diluted in RGBTw for 1 hour at room temperature. Following several washes with TBSTw, tissues sections were incubated with 4’,6-diamidino-2-phenylindole, dilactate (DAPI) to visualize cell nuclei and mounted in phosphate buffered saline containing 80% glycerol and 0.2% n-propyl gallate. Images were obtained using the Keyence BZ-X700 microscope (Keyence, Osaka, Japan). For primary and secondary antibody information see Table 1.

### Bulk population sequencing and analysis

Total RNA from 500K FACS-isolated basal epithelia, luminal epithelia, other epithelia, and stromal cell gates from 4 donor specimens was extracted using RNEasy micro columns (Qiagen). RNA quantity and quality were tested and samples were processed for RNA-Seq on a NextSeq 500 Sequencer (Illumina) in the UT Southwestern McDermott Center Next Generation Sequencing Core. The libraries were sequenced as stranded single-end 75 cycle reads. Analysis was done using the UT Southwestern Bioinformatics Core Facility RNA-Sequencing analysis workflow (https://git.biohpc.swmed.edu/BICF/Astrocyte/rnaseq). An average of 27 million sequencing reads per sample were aligned with HISAT2 to GRCh38^54^ at an average rate of 88%; duplicates are removed using SAMtools^55^; and counts are generated using FeatureCount^56^ using the annotations from Gencode V20^57^. Genes identified as Globins, rRNAs, and pseudogenes are removed. Differential expression analysis is performed using edgeR^58^, using a FC cutoff of 2 and adjusted FDR cutoff of 0.05. Pan-epithelial DEGs are an intersection of DEGs up in the different epithelial subpopulations (basal, luminal, and other) compared to stroma. Similarly, the stromal DEGs are the intersection of the DEGs which are up in stroma compared to the different epithelial subpopulations. The DEGs associated to each of the epithelial subpopulations are generated compared to stroma but are filtered to be unique for that subpopulation. Quantitative Set Analysis for Gene Expression (QuSAGE) was utilized to perform gene set enrichment-type analysis compared to publicly available prostate genesets^7,8,12^. Oudes *et al* published DEGs were used, but Zhang et al and Liu et al were calculated using limma (version 3.32.4)^59^ and a FDR cutoff of 0.05 (Benjamini-Hochberg corrected) of log 2 normCounts and log 2 FPKM, respectively.

### Single cell sequencing

Three young human prostate specimens were used for single cell sequencing. Single cell suspensions that were flow sorted for viability or gated for specific populations were loaded into the 10x Genomics Chromium Controller using the Chromium Single Cell 3’ Library and Gel Bead Kit v2 according to the manufacturer’s protocol. Briefly, 17,400 total cells of each sample were loaded on individual lanes of a Single Cell A Chip with appropriate reagents and run in the Chromium Controller to generate single cell gel bead-in-emulsions (GEMs) for sample and cell barcoding. Libraries were generated using 10x Genomics’ protocol. Libraries were pooled and submitted for sequencing on an Illumina NextSeq 500 in high output mode. 75 cycle flow-cells were used to sequence 26 cycles for read 1, 58 cycles read 2, and 8 cycles for the i7 index.

### Single cell sequencing data analysis

#### Three patient aggregate

Three patient specimens dissected into transition/central zone and peripheral zone. Each zone was sorted for viability before loading into the 10x Genomics chromium controller. The 10x Genomics’ analysis pipeline, cellranger (version 2.1.1) was first used to demultiplex and produce a gene-cell matrix. Bcl files were demultiplexed using their barcode-aware wrapper for bcl2fastq (version 2.17.1.14). Transcriptomes were aligned to GRCh38 using STAR (version 2.5.1b). Samples were then aggregated by downsampling to match their mean mapped reads per cell. Low quality cell barcodes were filtered out using 10x Genomics’ algorithm (high quality barcodes = total UMI count ≥ 10% of the 99^th^ percentile of the expected recovered cells). Supplementary Table 2 displays the sequencing metrics for each barcoded experiment.

Seurat (version 2.3.1), an R toolkit for single cell transcriptomics formed the basis of further analysis^13^ run on R version 3.4.1. Genes that were expressed in three cells or less were filtered out along with cells expressing fewer than 200 unique genes. Cell cycle state was predicted based on Seurat’s built in principal-component (PC) analysis. Briefly, cells were scored based on expression their expression of G2M and S phase genes^60^. Low quality cells and multiplets were excluded by removing cells with fewer than 500 unique genes and greater than 3,000 unique genes, as well as cells with greater than 10% of their transcriptome being mitochondrial genes. Data was then scaled to 10,000 and log transformed. Mitochondrial genes were then removed from further analysis. UMI counts were then scaled and variation due to differences in UMI/cell, percent mitochondrial genes, and cell cycle phase were regressed out of the data using a built-in Seurat function. Cells from the three patients were then subsetted and recombined using canonical correlation analysis (CCA) in order to align the clusters. The highest variable genes were found with an algorithm developed by Macosko *et al*^61^, and were defined as an average expression between 0.2 and 5 with a dispersion greater than 1. The intersection of these genes between the three patients were used to calculate 50 CCAs and the first 30 were aligned. These 30 aligned-CCAs were used for t-SNE visualization and clustering.

Cells were clustered using a graph-based clustering approach^13^. Briefly, cells were embedded in a KNN graph structure based on their Euclidean distance in PC space, with edge weights refined by shared overlap in their Jaccard distance. Different resolutions were generated based on a granularity input.

Highly stressed cells were predicted and removed by performing PC analysis on the cells’ expression of an MSigDB^16,17^ list of stress response genes (M10970) (Supplementary Table 3)^62^. The cells’ projection to PC1 was used as a “stress score”. To ensure this score is intuitive (stressed cells are more positive than unstressed), under the assumption that most of the cells are less stressed, the values are scaled such that the mean of the distribution is to the left of an expected normal distribution centered around zero. Highly stressed cells were chosen as 10% of the cells which had the top ‘stress score’. Clusters (from an over-clustered resolution of 1) were removed if at least 50% of the cells were identified as highly stressed. All remaining highly stressed cells were also removed and used to create a prostate-specific stress signature (see Supplementary Figure 3 and Supplementary Table 3) by calculating DEGs compared to lowly-stressed cells with a Wilcoxon rank sum test on genes present in at least 50% of either group which are at least five-fold enriched in the stressed cells and a maximum Bonferroni corrected p-value alpha of 0.05. Remaining cells were then re-clustered and new t-SNE plots were generated.

For cluster cell lineage (epithelial or stroma), the expression of genes in each cluster was correlated using QuSAGE (version 2.6.0) to 5-fold change differentially expressed genes (5FC-DEG) of pan-epithelial and stromal transcriptomes obtained from bulk population RNA-Seq. The cells were clustered at a resolution of 0.2 and QuSAGE was performed on a random subset of cells from each cluster, sampled to the smallest cluster. All identities were assigned from correlation as the highest positive enrichment score. Each lineage was independently sub-clustered, and t-SNE recalculated for cell-type identification. Epithelial clusters were correlated to 5FC-DEGs from population RNA-Seq of basal epithelia, luminal epithelia, and “other” epithelia. Stromal clusters were correlated to a subset of GeneOntology biological process gene sets related to known stromal cell types. The gene sets used were GO_ENDOTHELIAL_CELL_DIFFERENTIATION, GO_SMOOTH_MUSCLE_CELL_ DIFFERENTIATION, GO_REGULATION_OF_FIBROBLAST_PROLIFERATION, and GO_LEUKOCYTE_ACTIVATION (see Supplementary Figure 4). The epithelial and stromal QuSAGE analysis, similar to the lineage analysis, was performed on random subsets of cells in each cluster, sampled to the smallest cluster in the specific analysis.

Rare neuroendocrine (NE) cells were identified from the epithelial cells using an initial NE gene set from Table 1 of Vaschenko et al^63^ and a PC-based method was used for stress identification, with the only difference being NE cells were chosen as the 0.1% of epithelial cells with the highest ‘NE score’ (see Supplementary Figure 5). A revised NE DEG list based on the difference between identified NE cells and the other epithelial cells is found in Supplementary Table 3. The DEGs were determined with a Wilcoxon rank sum test on genes present in at least 1% of either group with a maximum Bonferroni corrected p-value alpha of 0.05.

Cluster/cell identities were aggregated and DEGs for the identified cell types were then determined using a Wilcoxon rank sum test on genes present in at least 25% of either the population of interest or all other cells in that lineage which are at least two-fold enriched in the population of interest and a maximum Bonferroni corrected p-value alpha of 0.05.

Monocle^26^ was performed on sub-setted epithelial cells using to predict possible differentiation trajectory in pseudotime.

#### FACS

Cells from basal, luminal, ‘other’ epithelia, and fibromuscular were sorted out separately and single cell RNA-sequencing was performed in a similar fashion to the three patient aggregate. These samples were analyzed to demonstrate the rate of contamination of the FACS gates. Instead of running CCA (as all of the samples are from the same patient and didn’t need alignment), PCA was used conducted on the highest variable genes (same method as above). The PC’s representing the top 85% of the cumulative variation of 50 calculated PC’s was used for clustering and t-SNE calculation. Also, the novel DEG lists created above from the three patient aggregate experiment were used for cell type identification using QuSAGE, and stress and NE identification using PCA. This experiment was conducted on a second patient using a novel FACS panel predicted from the three patient aggregate DEG lists. The populations analyzed in the second patient was, basal epithelia, luminal epithelia, ‘other’ epithelia, ‘double-negative’ epithelia, fibroblasts, and smooth-muscle. Table 1 displays information on antibodies used for flow cytometry.

#### Downsample experiment

One library was sequenced to deeper (382,925,951 reads, resulting in an average of 75,082 reads per cell), and the reads were randomly sampled to determine the relationship between cluster integrity and cell identity with read depth. Each of the sampled fastq sets were run through the pipeline the same as the FACS samples. The cell type identities were subjected to normalized mutual information (NMI) analysis, to quantify the mutual dependence of each sample to the un-sampled data (ground truth). The NMI was fitted to the mean reads per cell using locally weighted scatterplot smoothing. The minimum reads per cell that is required to produce a NMI of 0.9 was calculated from the model.

### qPCR analysis

For quantitative real-time PCR (qPCR), RNA was extracted with Trizol (Ambion) from 200-500K flow cytometry-isolated cells. RNA was reverse transcribed into cDNA using RT^2^ First Strand Kit (Qiagen, Valencia, CA). qPCR was performed using IQ SYBR Green Supermix (BioRad, Hercules, CA) and results were analyzed using BioRad CFX manager software. All results were calculated using ΔΔCt analysis and normalized to RPL27 expression. Statistical significance was calculated by t test using Graphpad Prism software (version 7.0d). Primer sequences are listed in Table 1.

### Data and software availability

To increase rigor, reproducibility and transparency, raw image files (including duplicates not displayed here), raw FACS data and RNA-seq data generated as part of this study were deposited into the GUDMAP consortium database and are fully accessible at: https://doi.org/10.25548/W-R8CM. R code used to produce all the single cell RNS-seq analysis can be found at https://git.biohpc.swmed.edu/StrandLab/sc-TissueMapper_Pr.git. Analyzed data from the three patient aggregate single cell RNA-seq experiment can be found at http://strandlab.net/analysis.php, where gene expression can be investigated in the cell type clusters identified in this study.

### Accession numbers

The bulk and single cell RNA-seq data from normal prostate tissues were deposited into the GEO SuperSeries GSE120716.

## Acknowledgments

We thank the families of organ donors at the Southwest Transplant Alliance for their commitment to basic science research. Financial support came from K01 DK098277, R03 DK110497, R01 DK115477, and U54DK104310 pilot program award (D.W.S.); U54DK104310, U01DK110807 and R01DK099328 (C.M.V); V.S.M. and J.L. are supported by a grant from CPRIT (RP150596); Cancer Prevention Research Institute of Texas (CPRIT) (RR140023 to G.C.H.), NIH (DP2GM128203 to G.C.H.), the Department of Defense (PR172060 to G.C.H.), the Welch Foundation (I-1926-20170325 to G.C.H.), and the Green Center for Reproductive Biology; VA North Texas Health Care System New Investigator Program award #16-001 (J.C.G.), and the generous donations of the Smith, Penson, and Harris families to the UTSW Department of Urology (C.G.R.).

## Author contributions

Conceptualization, D.W.S.; Methodology, D.W.S, G.H.H., G.C.H., and C.M.V.; Software, G.H.H, V.S.M. and J.L.; Validation, D.W.S. and C.M.V.; Formal Analysis, D.W.S., G.H.H., and C.M.V.; Investigation, D.W.S., G.H.H., A.M., D.B.J., R.M., J.C.R., and R.C.H.; Resources, D.W.S, G.V.R., M.P.M., J.C.G., R.C.H., J.C.R., and C.M.V.; Data Curation, G.H.H. and D.B.J.; Writing – Original Draft, D.W.S., G.H.H.; Writing – Review and Editing, D.W.S., G.H.H., G.C.H., V.S.M., J.L., G.V.R, and C.M.V.; Visualization, D.W.S.; Supervision, D.W.S. and C.M.V.; Project Administration, D.W.S.; Funding Acquisition, D.W.S., J.C.G., G.V.R., C.M.V., and C.G.R.

## Supplemental Information titles and legends

**Supplementary Figure 1.**
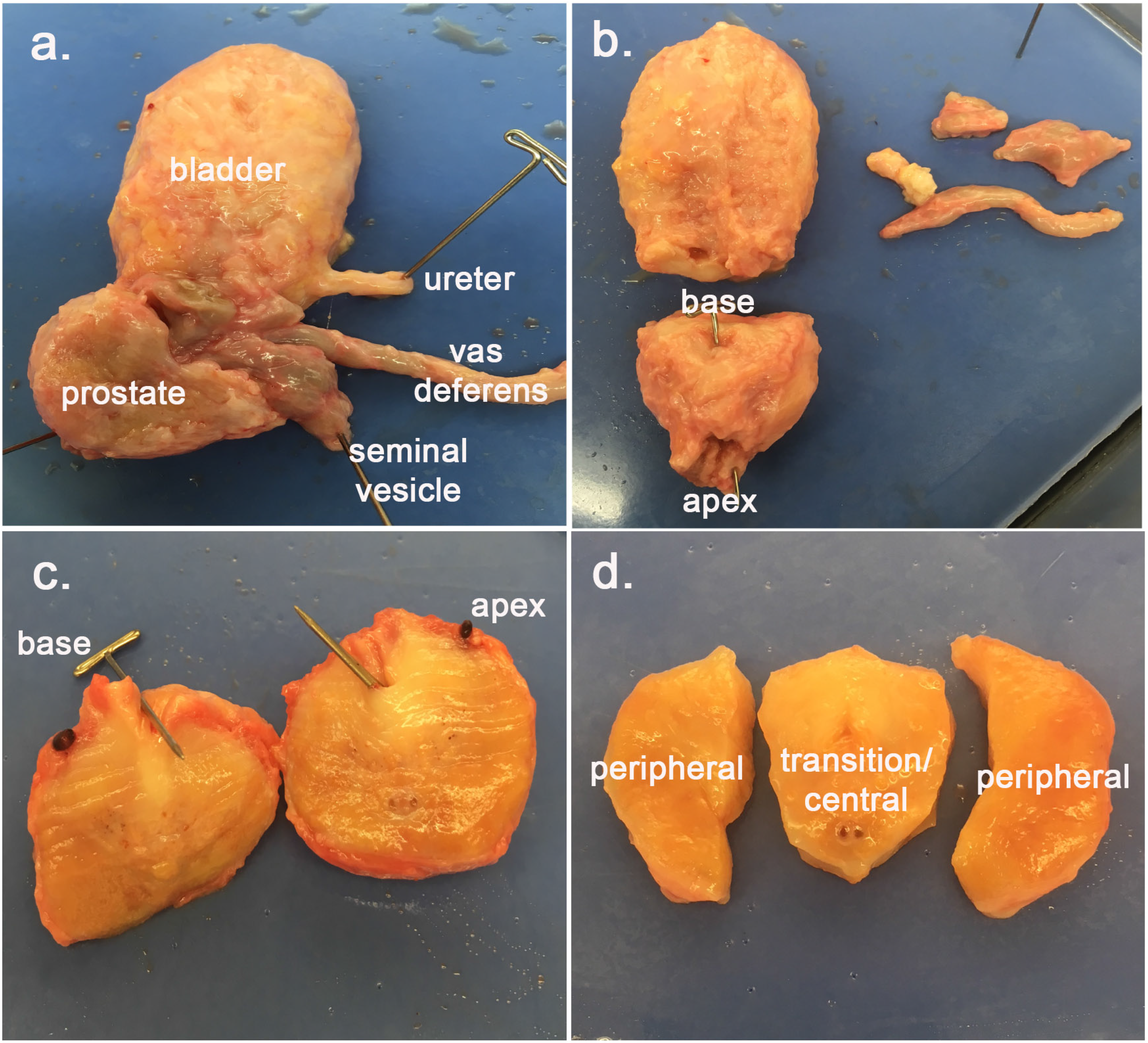
Dissection of human prostate. (a) A cystoprostatectomy is performed on young organ donors after transplantable organ harvest. (b) The bladder, vas deferens, and seminal vesicles are separated from the prostate. (c) A coronal section is made for generation whole mounts and for (d) dissecting the peri-urethral transition zone.

**Supplementary Figure 2.**
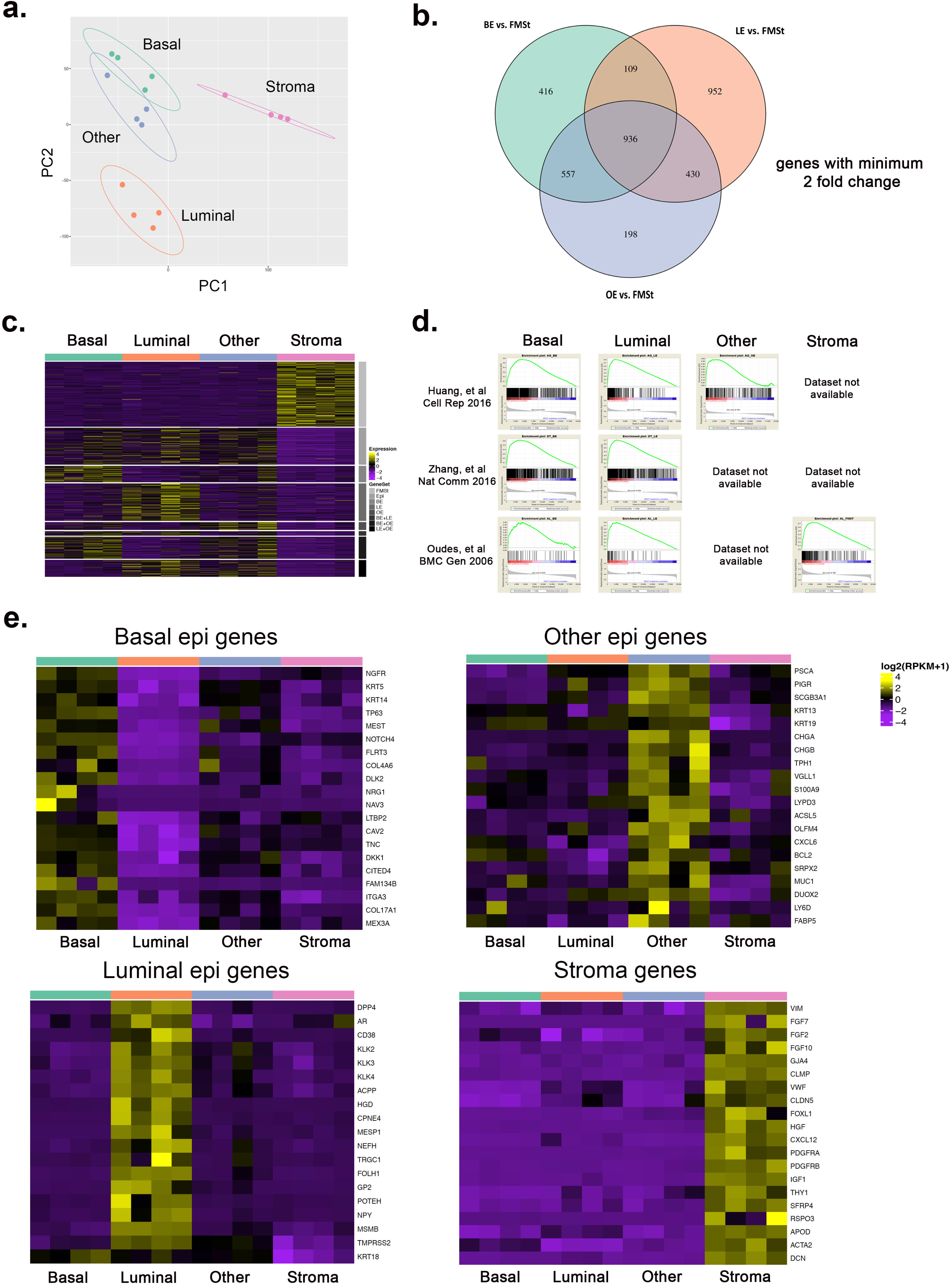
Bulk sequencing of normal human prostate cell types isolated by flow cytometry. (a-c) Principal component analysis, Venn diagram, and heat map of DEGs from basal epithelia (CD45^-^/CD31^-^/CD326^+^/CD271^+^/CD26^-^), luminal epithelia (CD45^-^/CD31^-^/CD326^+^/CD271^+^/CD26^+^), ‘other’ epithelia (CD45^-^/CD31^-^/CD326^+^/CD271^-^/CD26^-^), and stroma (CD45^-^/CD31^-^/CD326^-^). (d) GSEA of flow cytometry-isolated, bulk sequenced cell types from young normal prostate *vs*. enriched cell populations from 3 published data sets. BE, Basal epithelia; LE, Luminal epithelia; FMSt, Fibromuscular Stroma. (e) Heat maps of DEGs from Basal, Luminal, and Other epithelial gates as well as Fibromuscular stroma with 20 genes highlighted.

**Supplementary Figure 3.**
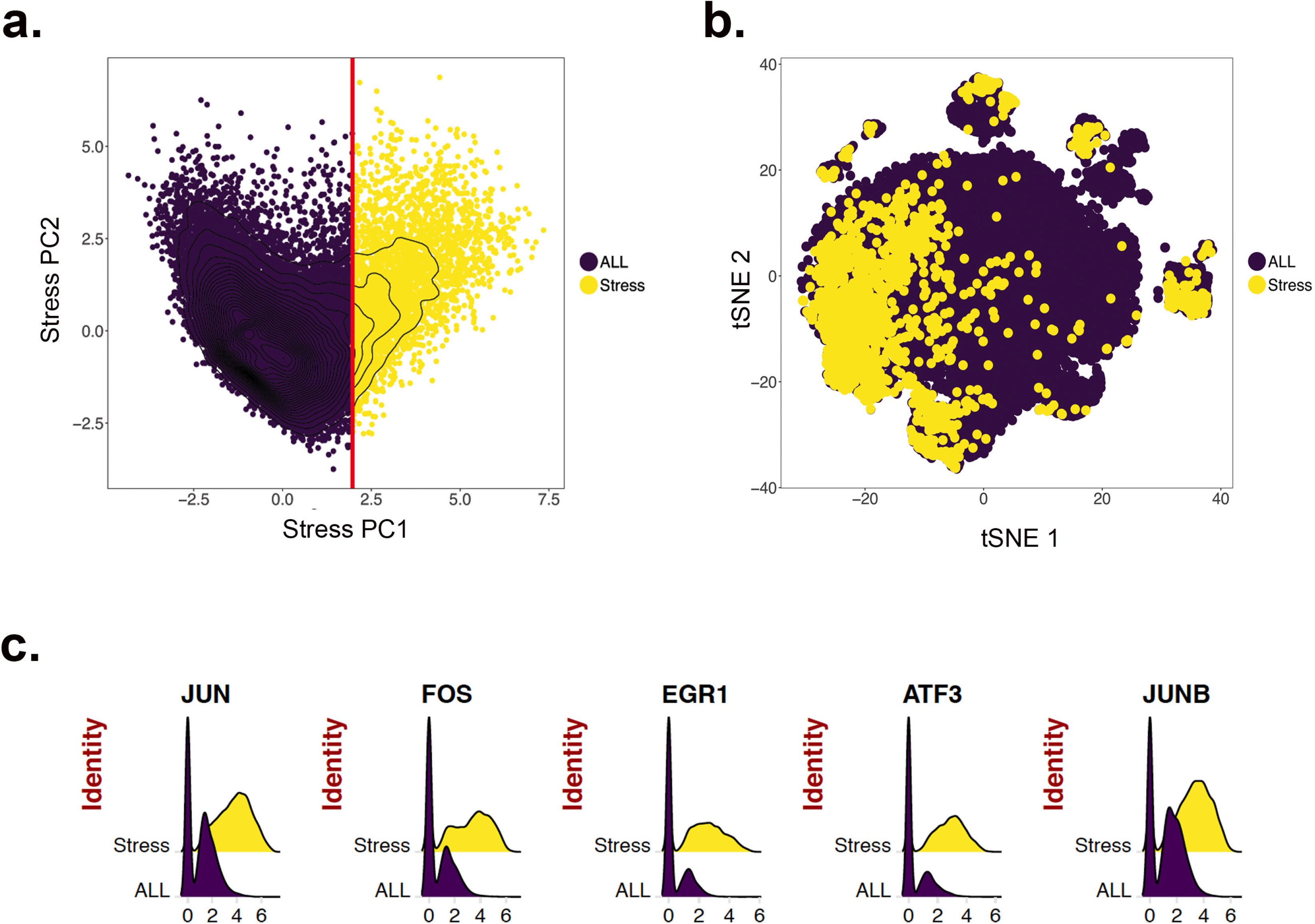
Bioinformatics approach to identification and removal of stressed cells. (a) A signature of cell stress from MSigDB was used to create a pseudogene for principal component analysis. (b) Stressed cells were identified in every cluster. (c) Ridge plots of the top 5 stress signature genes in stressed cells *vs*. unstressed cells (ALL). cells.

**Supplementary Figure 4.**
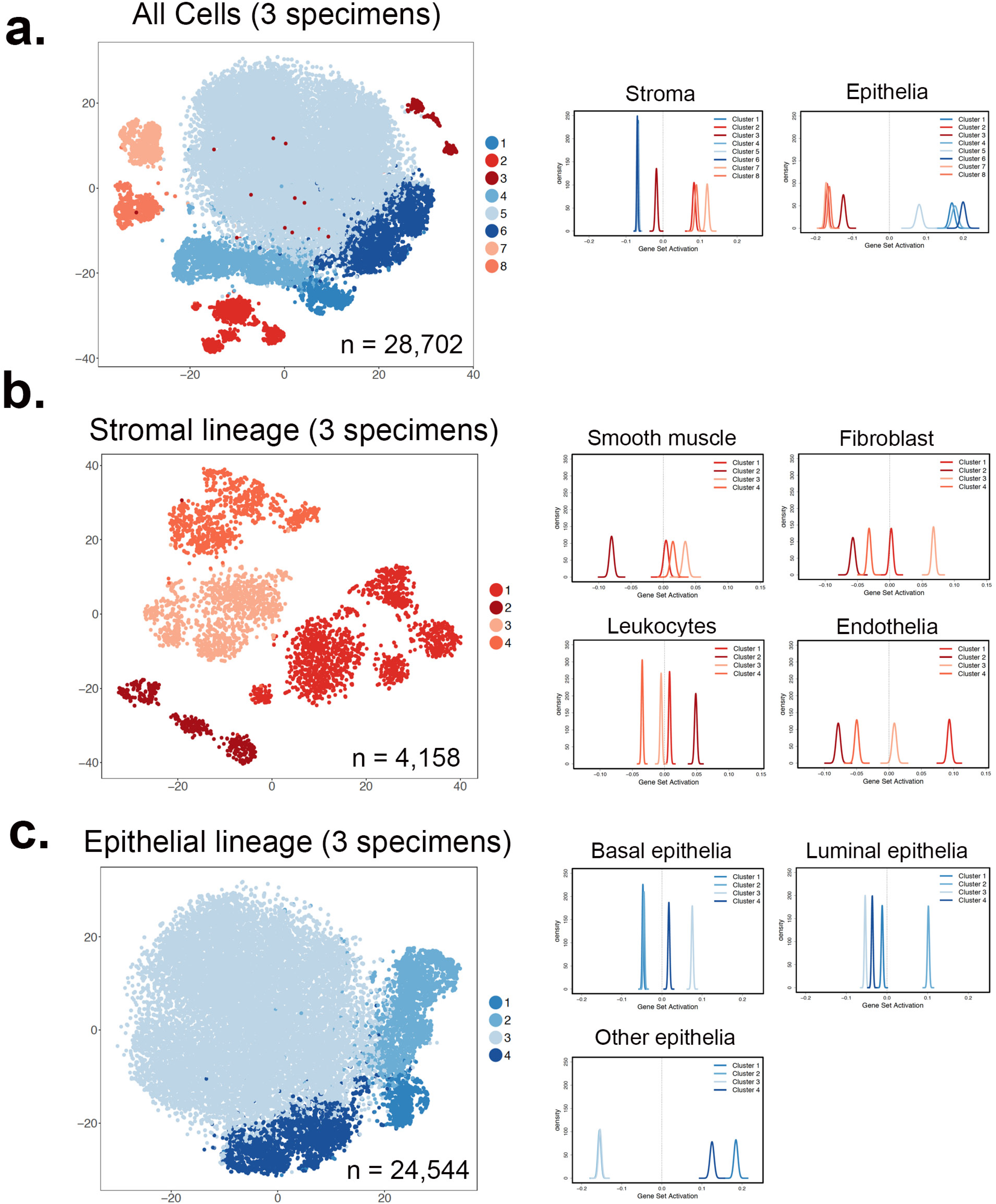
Method of cell cluster identification. (a) Bulk sequencing of CD45^-^/CD31^-^/CD326^-^ prostate stroma and CD326^+^ epithelia was used to identify stromal and epithelial cell lineages for sub-clustering. (b) Gene ontology terms for smooth muscle, fibroblast, endothelia, and leukocytes were used in GSEA to identify stromal sub-clusters. (c) Bulk sequencing of CD271^+^ basal, CD26^+^ luminal, and CD271^-^/CD26^-^ ‘other’ epithelia was used to identify epithelial clusters.

**Supplementary Figure 5.**
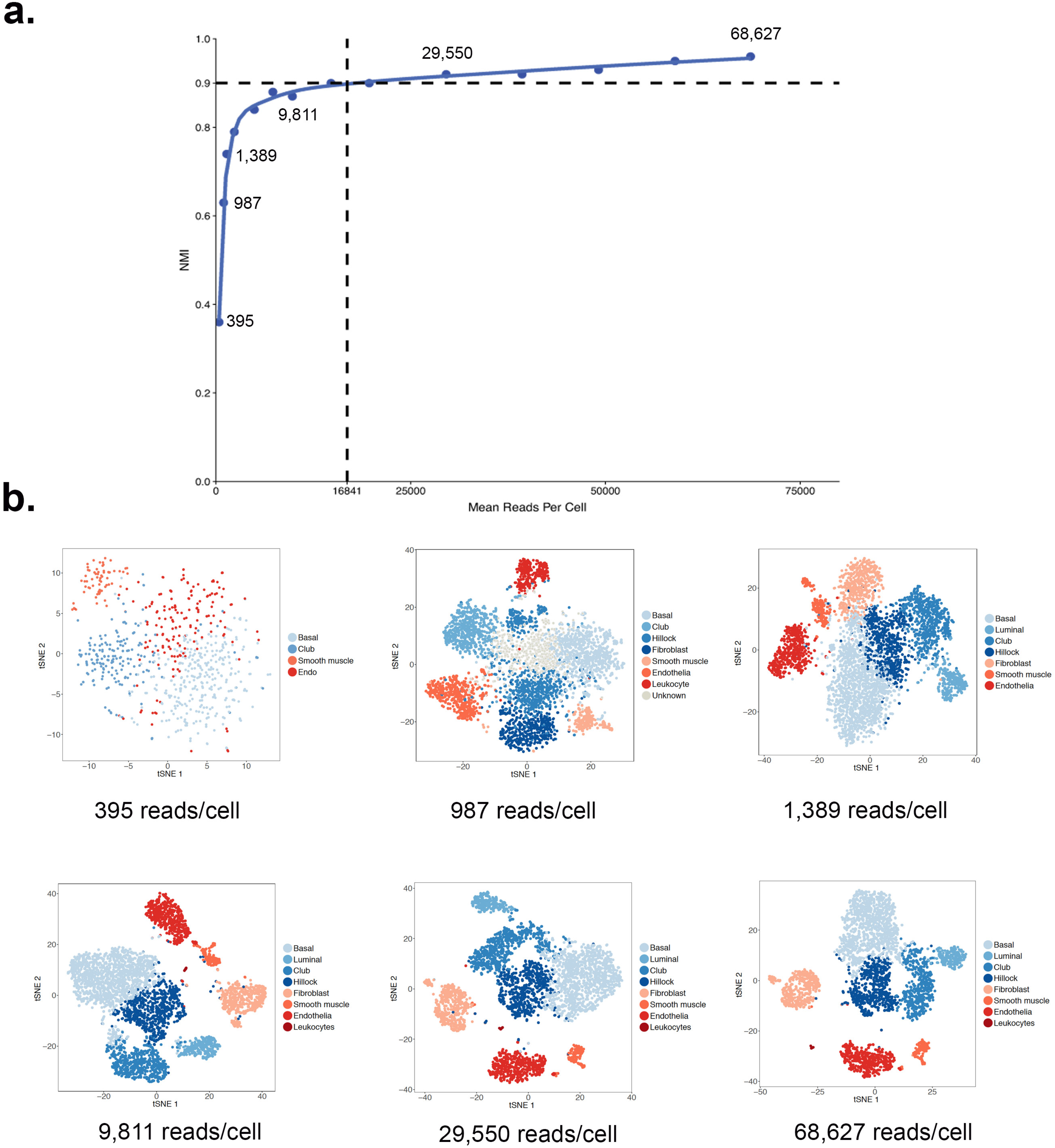
Downsampling experiment to determine effect of read depth on cluster identification. (a) Mean read depth per cell plotted against the normalized mutual information (NMI) compared to the unsampled data. Dotted line represents the minimum average read depth per cell required to maintain a NMI of 0.9. (b) tSNE plots of identified clusters from representative read depth per cell experiments.

**Supplementary Figure 6.**
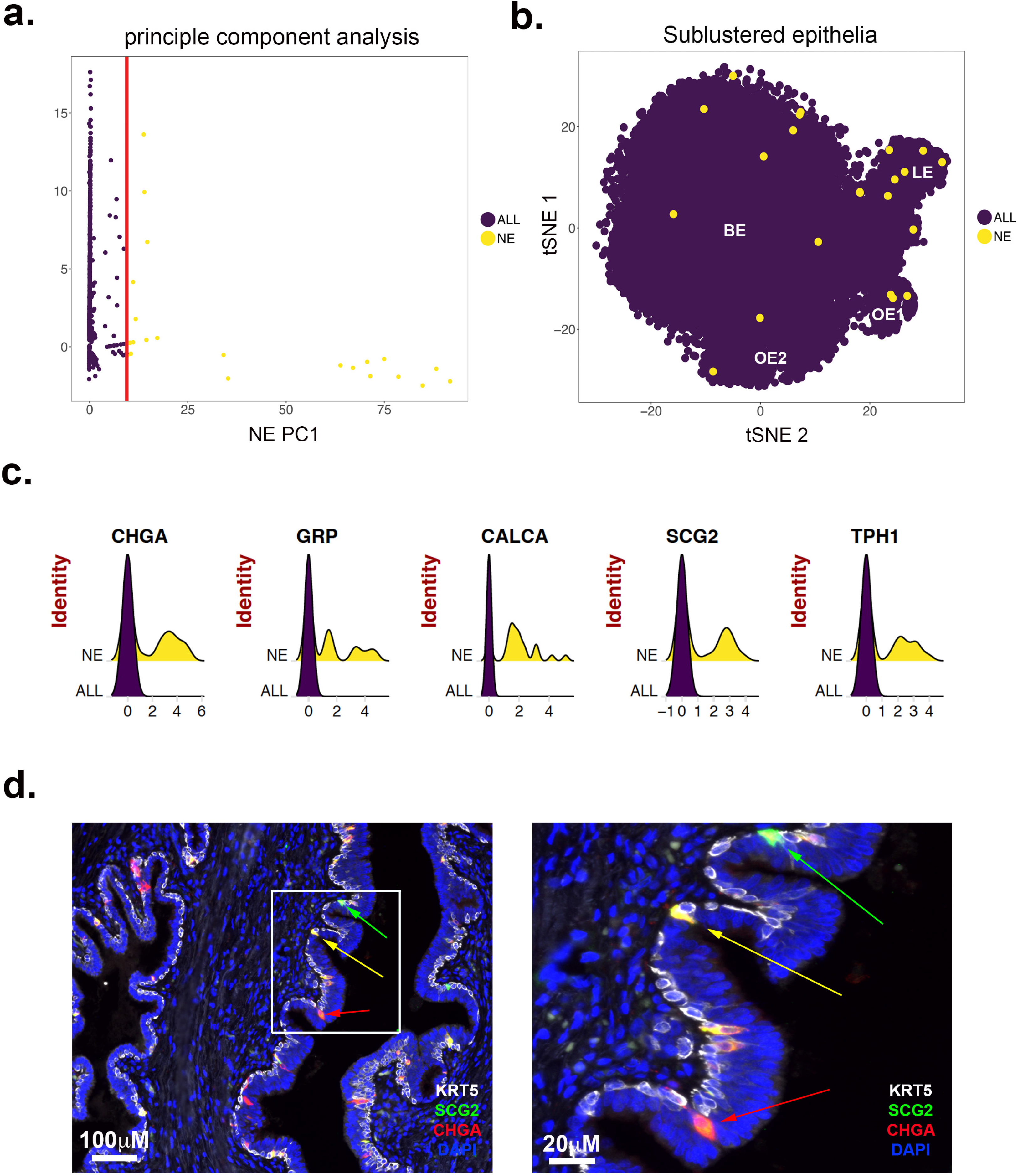
Supervised identification of neuroendocrine epithelia (NE). (a) A signature of NE epithelia (Supplementary Table 3) was used to create a pseudogene for principal component analysis, which led to the detection of 25 NE cells in the scRNA-seq dataset of three donor specimens. (b) NE epithelia were found mainly within the luminal cell cluster, but also in other clusters. (c) Rigde plots of the top 5 DEGs derived from comparing NE cell transcriptomes to non-NE epithelial transcriptomes. (d) Immunofluorescence for cytokeratin 5 (basal marker in white), CHGA (classic NE marker in red) and SCG2 (novel NE marker in green) showed both overlapping (yellow) and non-overlapping expression (red or green).

**Supplementary Table 1.** List of significant DEGs generated by bulk RNA sequencing of FACS-isolated human prostate cell types.

**Supplementary Table 2.** Sequencing metrics for human prostate scRNA-seq experiments.

**Supplementary Table 3.** List of DEGs exclusive to each cell population.

**Supplementary Table 4.** QuSAGE analysis of the C2 dataset of MSigDB for each cell cluster.

